# Advanced deep-tissue imaging and manipulation enabled by biliverdin reductase knockout

**DOI:** 10.1101/2024.10.18.619161

**Authors:** Ludmila A. Kasatkina, Chenshuo Ma, Huaxin Sheng, Matthew Lowerison, Luca Menozzi, Mikhail Baloban, Yuqi Tang, Yirui Xu, Lucas Humayun, Tri Vu, Pengfei Song, Junjie Yao, Vladislav V. Verkhusha

**Affiliations:** Department of Genetics and Gruss-Lipper Biophotonics Center, Albert Einstein College of Medicine, Bronx, NY 10461, USA; Department of Biomedical Engineering, School of Engineering, Duke University, Durham, NC 27708, USA; Department of Anesthesiology, School of Medicine, Duke University, Durham, NC 27708, USA; Department of Electrical and Computer Engineering, University of Illinois Urbana- Champaign, Urbana, IL 61801 USA; Medicum, Faculty of Medicine, University of Helsinki, Helsinki 00290, Finland

## Abstract

We developed near-infrared (NIR) photoacoustic and fluorescence probes, as well as optogenetic tools from bacteriophytochromes, and enhanced their performance using biliverdin reductase-A knock-out model (Blvra-/-). Blvra-/- elevates endogenous heme-derived biliverdin chromophore for bacteriophytochrome-derived NIR constructs. Consequently, light-controlled transcription with IsPadC-based optogenetic tool improved up to 25-fold compared to wild-type cells, with 100-fold activation in Blvra-/- neurons. *In vivo*, light-induced insulin production in Blvra-/- reduced blood glucose in diabetes by ∼60%, indicating high potential for optogenetic therapy. Using 3D photoacoustic, ultrasound, and two-photon fluorescence imaging, we overcame depth limitations of recording NIR probes. We achieved simultaneous photoacoustic imaging of DrBphP in neurons and super-resolution ultrasound localization microscopy of blood vessels ∼7 mm deep in the brain, with intact scalp and skull. Two-photon microscopy provided cell-level resolution of miRFP720-expressing neurons ∼2.2 mm deep. Blvra-/- significantly enhances efficacy of biliverdin-dependent NIR systems, making it promising platform for interrogation and manipulation of biological processes.

## INTRODUCTION

Optogenetic manipulation and molecular imaging *in vivo* are essential approaches to control and monitor cellular events. *In vivo* optogenetics and molecular imaging can greatly benefit from optogenetic tools (OTs)^1-3^ and imaging probes^4-6^ that operate in the near-infrared (NIR: 650-900 nm) spectral range. Due to low tissue absorbance, low autofluorescence and reduced light-scattering^7^ in the NIR range, these systems enable deep-tissue penetration, higher resolution, and sufficient light power for optogenetic manipulation.

Almost all currently available NIR OTs, NIR fluorescent proteins (FPs)^8, 9^, and NIR photoacoustic (PAT) probes have been engineered from soluble bacterial phytochrome photoreceptors (BphPs)^2, 9^. Nevertheless, most widely-used optical imaging technologies, such as confocal, two-photon, or optical coherence tomography, all rely on the (quasi)ballistic photons and thus lack penetration depths beyond the first 1-2 millimeters from the skin surface^7, 10^, which fundamentally hinders their applications *in vivo*. By contrast, strong NIR absorption of BphPs allows their use as molecular probes for photoacoustic tomography (PAT). PAT detects light-induced ultrasound waves and has intrinsic sensitivity to the optical absorption contrast of biomolecules. Because the biological tissues usually have much weaker attenuation for acoustic waves than light, the imaging depths attained by PAT are far beyond that attainable by other purely optical imaging approaches^11-13^. We and others previously demonstrated that PAT can take full advantage of the strong NIR light absorption of BphP and achieve high-resolution (<0.5 mm) imaging of BphP-expressing tumors or organs at centimeter-level depths in soft tissues^11, 12, 14-18^. Importantly, the reversible photoswitching capability of BphPs between two absorption states enabled the differential detection in PAT, also called reversible-switching PAT (RS-PAT), which effectively suppressed the non-switching background signals majorly from blood and improved the molecular detection sensitivity by three orders of magnitude when compared with PAT using genetically-encoded non-switching probes^11, 12^.

BphPs and the BphP-derived proteins use biliverdin IXa (BV) as a NIR chromophore^19^,^20^. In mammals, including humans, these probes can incorporate endogenously produced BV into its chromophore-binding pockets, thus forming holoforms^21^. BV is an important cytoprotective and anti-inflammatory molecule formed in all tissues as the product of heme catabolism. A special role in BV formation *in vivo* is assigned to the mononuclear phagocyte system^22^ of the liver, spleen and lymph nodes, where senescent red blood cells undergo clearance^23^ (**Supplementary** Fig. 1a). Sequential enzymatic steps, catalyzed by heme oxygenase and BV reductase, convert heme into BV and bilirubin, respectively (**Supplementary** Fig. 1b). BV reductase converts BV to bilirubin via the reduction of a double-bond between the second and third pyrrole ring into a single-bond.

However, BV concentration varies greatly among tissues and cell types, being quite low in some organs like the brain, and, consequently, the amount of the formed functional holoform of the BphP-based NIR constructs in these tissues is low, resulting in an inability to do optogenetic activation in these tissues. This low amount of BV is the major obstacle to using the NIR constructs *in vivo*, and because all of them are BphP-derived, this significantly limits the optogenetic applications in deep tissues. This is also the most important reason for the lack of PAT applications in deep brains using BphP-derived molecular probes. It is particularly challenging for PAT to image BphP-labeled neurons due to the low BV level in the brain tissues.

To overcome this problem, several strategies were proposed. First, since low BphPs expression level favors its complete assembling with endogenous BV into a holoform^3, 24^ the generation of a transgenic mouse encoding only two copies of the *RpBphP1* BphP gene per cell demonstrated its advantage for *in vivo* applications^14^. Another strategy to increase the amount of the formed holoform *in vivo* is to either increase the production of BV or inhibit its conversion into bilirubin. We hypothesized that a much better way to have the higher BV level in tissues and, consequently the higher amount of the holoform of BphP-derived probes, would be to use BV reductase A (Blvra) knockout (termed Blvra-/-) animals. Blvra is the main isoform among just two BV reductases in mice and humans, each encoded by its own gene.

Here, we study the performance of iLight OT to drive gene expression in the case of Blvra knock-out in mice. iLight is based on the photosensory core module (PCM) of IsPadC BphP from *Idiomarina sp*. ^25^ and is coupled with the Gal4-UAS system, allowing efficient NIR light-controlled gene transcription in cells and *in vivo*. We first evaluated the activation of iLight in the primary cells isolated from Blvra-/- mouse. We next demonstrated the light-dependent correction of diabetes in Blvra-/- mice but not in wild-type (WT, i.e., Blvra+/+) animals. Moreover, by providing more efficient photoswitching and stronger fluorescence emission, the detection sensitivity of PAT and fluorescence imaging (both one-photon and two-photon fluorescence) of the BphP-derived molecular probes can be substantially improved by using the Blvra-/- model. Particularly, we developed a three-dimensional (3D) photoacoustic and ultrasound localization micrsocopy system (termed 3D-PAULM), which is capable of simultaneous PAT of BphP-derived probes and ultrasound localization microscopy (termed ULM) of the vasculature. Combining PAT and ULM into a single system can provide a more comprehensive visualization of the vasculature structures and molecular signatures, which is not available in our previous PAT systems. Applying 3D-PAULM on the Blvra-/- mice provided superior molecular imaging in various deep tissues *in vivo*, including brain, liver and spleen, as well as breast tumors expressing DrBphP-PCM. Furthermore, the one-photon and two-photon fluorescence level of BphP-derived probes in the deep tissues of Blvra-/- mice was also elevated.

## RESULTS

### Establishing mouse models

As our colony was established from heterozygous Blvra-/+ mice, for F_1_ progeny we used genotyping to detect breeders and experimental animals with double knock-out *Blvra* gene. PCR genotyping confirmed the inversion in the *Blvra* (**Supplementary** Fig. 2a) in all animals selected for further experiments (**Supplementary** Fig. 2b).

To perform PAT imaging of genomically encoded *R. palustris* RpBphP1 (below denoted as BphP1) bacterial phytochrome, we generated homozygous loxP-BphP1 mice with double knock-out of *Blvra* (termed “loxP-BphP1 Blvra-/-“). For this, loxP-BphP1 mice^14^, bearing the loxP-EGFP-loxP-BphP1-mCherry-TetR insert in *ROSA26* locus, were crossed to Blvra-/- mice. Following sequential breeding steps (**Supplementary** Fig. 3a), the progeny was genotyped using the primers for EGFP and intact *ROSA26* locus, as well as primers for WT and *Blvra* knockout allele. (**Supplementary** Fig. 3b). PCR genotyping was reconfirmed by *in vivo* imaging of EGFP fluorescence (**Supplementary** Fig. 3c). The loxP-BphP1 Blvra-/- mice have substantially more greenish gallbladders when compared with the loxP-BphP1 Blvra+/+, confirming the elevated BV level (**Supplementary** Fig. 3d). Double-homozygous loxP-BphP1 Blvra-/- mice were further crossed to each other to maintain the colony. According to the studies on the Blvra-/- mouse strains developed and characterized by several research groups^26-29^ there were no gross phenotype abnormalities observed in mice with knock-out of Blvra gene. Bvra deficiency had no gross effect on liver or kidney function^26^. Blvra-/- offspring were born without detectable anomaly at least until the 11^th^ generation^28^. In our observations during colony maintenance of Blvra-/- mice and loxP-BphP1 Blvra-/- mice for 2 years no changes were observed in mouse health, fertility, breeding, litter size, development, behavior, and lifespan.

### Optogenetic activation of gene expression in isolated primary cells

We analyzed the performance of iLight in several primary cell types from Blvra-/- mice, such as cortical neurons, lung endothelial cells, and skin fibroblasts, and compared it with that in WT primary cells. For this, we studied the iLight-induced transcriptional activation of luciferase reporter AkaLuc^30^. AkaLuc was cloned downstream of 12× UAS (denoted as G12 through the text) with TATA-box minimal promoter. Leakage is a known phenomenon for reporter genes and should be used as a control to correctly evaluate the light-induced transcription and to estimate the transcription activation with iLight (if any) in the darkness. Therefore, where necessary, we equilibrated the expression of the AkaLuc reporter gene with AAV encoding iGECInano sensor (AAVeq) earlier developed by us, and transduced cells using the same viral particles ratio (AAV iGECInano: AAV G12-AkaLuc, 1:3) and multiplicity of infection (10^5^) as was used for samples transduced with AAV iLight and AAV G12-AkaLuc. In addition, to estimate the impact of exogenous BV in both WT and Blvra-/- genotypes, separate sets of cells were supplemented with BV immediately after AAV transduction. Plates were kept in darkness or illuminated using a 660/15 nm LED array (light power density of 0.4 mW cm^-2^) for 48 h (**Fig. 1a)**. The AkaLuc expression level was measured by bioluminescence imaging (**Fig. 1b**). The iLight performance was determined as the fold increase of the average bioluminescence radiance in illuminated cells compared to cells kept in darkness (light-to-dark ratio, **Fig. 1c-e**).

**Figure 1.**
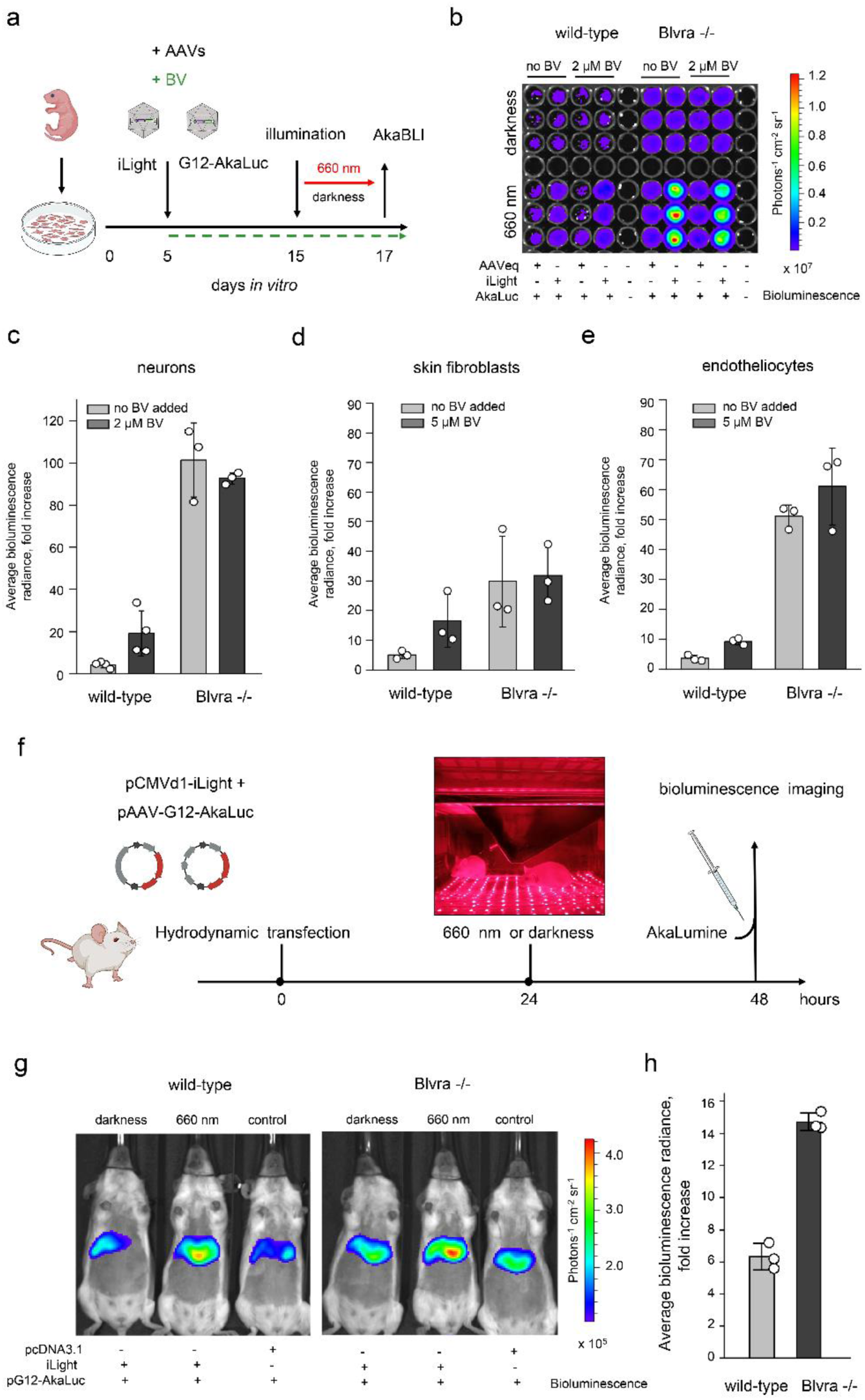
Optogenetic activation of gene transcription in primary cells and *in vivo*. **(a)** Experimental design. Primary cells were isolated from P0-P1 postnatal WT and Blvra-/- mice. Five days after isolation cells were transduced with AAV8 delivering iLight (CMVd1-NLS-Gal4-iLight-VP16) and AkaLuc reporter gene (G12-AkaLuc) (1:3, multiplicity of infection of 10^5^). Where necessary cells were supplemented with exogenous BV (green dashed line). **(b)** For AkaLuc bioluminescence imaging cells were transferred into the black 96-well plate and supplemented with 250 µM AkaLumine. Detection was performed using the IVIS Spectrum instrument. The rows represent 3 technical replicates of one independent experiment among 4 (WT cells) or 3 (Blvra-/- cells) performed in total on primary neurons. **(c-e)** iLight-mediated AkaLuc expression in primary WT and Blvra-/- cells. Performance of the OT was calculated as the fold increase of the average bioluminescence radiance in illuminated cells and cells kept in darkness (light-to-dark ratio) for neurons (c), skin fibroblasts (d), and endothelial cells (e). (**f**) Experimental design for optogenetic transcriptional activation in WT and Blvra-/-mice. Light-induced expression of AkaLuc in the liver was detected after hydrodynamic co-transfection with pCMVd1-NLS-Gal4-iLight-VP16 and pAAV-G12-AkaLuc. After illumination of animals with 660/15 nm LED array (3.5 mW cm^-2^) for 24 h the bioluminescence in the liver region was recorded. Animals were injected with AkaLumine (0.075 nmol/g of body weight) 15 min before imaging. **(g, h)** *In vivo* bioluminescence imaging of the iLight-induced expression of AkaLuc in the liver (g). The average bioluminescence radiance was calculated in the liver region and the light-to-dark ratio for transcriptional activation of AkaLuc (h) was calculated after subtraction of the background signal produced by the reporter plasmid only (pAAV-G12-AkaLuc with empty pcDNA3.1+). Data are presented as mean values +/-SD (*n*=3; independent transfection experiments).

Primary Blvra-/- neurons demonstrated significant enhancement in iLight performance, from 4.5-fold light-to-dark AkaLuc expression ratio in WT cells to nearly 100-fold ratio in Blvra-/- cells (**Fig. 1c**). Supplementation of primary neurons with exogenous BV resulted in a 19.2-fold light-to-dark ratio in WT cell (i.e. ∼4-fold improvement over WT neurons without external BV) whereas Blvra-/- cells demonstrated no change in iLight performance (**Fig. 1c**). In skin fibroblasts iLight performance was ∼6-fold better in Blvra-/- cells than in WT cells (5.1-fold vs. 30-fold light-to-dark AkaLuc expression), and supplementation with BV enhanced the light-to-dark ratio of AkaLuc production by ∼16.6-fold in WT cells, but did not affect the iLight performance in Blvra-/-cells (**Fig. 1d)**. In WT endothelial cells optogenetic activation of AkaLuc production was comparable to that of skin fibroblasts, i.e., a 3.5-fold light-to-dark ratio without and a 9.2-fold with exogenous BV (**Fig. 1e)**. iLight performance was ∼14.5-fold better in Blvra-/- in endothelial cells: without exogeneous BV the light-to-dark ratio was 51-fold, and slightly increased upon BV addition.

Thus, the iLight performance in Blvra-/-cells enhanced up to 25-fold over WT cells, and external BV almost did not affect the light-to-dark ratio of AkaLuc reporter expression, suggesting that endogenous BV level in Blvra-/- cells was sufficiently high to fully convert iLight apoform in the holoform.

### Comparison of iLight-mediated gene expression in WT and Blvra-/- mice

We next evaluated iLight performance *in vivo* in the liver of WT and Blvra-/- animals. Mice of the same genotype, age, sex and preferably littermates were used in the experiments to minimize the variables affecting the results. In line with permanent animal identification (eartags) we used short-term tail marking for convenience. The liver and spleen represent major organs of the mononuclear phagocyte system where senescent erythrocytes are deposited and heme is converted to BV and bilirubin for excretion. This feature distinguishes the liver and spleen in live animals from isolated hepatocytes and many other primary cells *in vitro*. To determine whether *Blvra* knock-out can further improve optogenetic manipulation in the liver of live mice, we hydrodynamically co-injected mice via tail vein with pCMVd1-NLS-Gal4-iLight-VP16 and pG12-AkaLuc plasmids in 1:3 ratio (**Fig. 1f**) and measured bioluminescence after 24 h of illumination with 660/15 nm LED array (**Fig. 1g**). Light power density for illumination of live mice was 9 times higher compared to that for cells in culture (3.5 mW cm^-2^). The same LED array, shown in **Fig. 1f**, was used for illumination *in vitro* and *in vivo*. To evaluate possible leakage of the AkaLuc reporter we used mice co-injected with pG12-AkaLuc and pcDNA3.1+ plasmids (“control” in **Fig. 1g**). Although absolute signals may slightly vary from one experiment to another, the difference between the bioluminescence evoked by AkaLuc reporter leakage and bioluminescence produced by the full-component OT (iLight + AkaLuc reporter) in the darkness was small and this tendency was preserved among the experiments. The bioluminescence produced by AkaLuc reporter-only control was subtracted from the values produced by the full-component OT (in the darkness and upon illumination) and then the light-to-dark ratio of optogenetic activation was calculated. Based on three independent optogenetic experiments the light-to-dark ratio for iLight-mediated AkaLuc expression was 2.3-fold higher in the livers of Blvra-/- mice compared to WT animals **(Fig. 1h)**, suggesting that the *Blvra* knock-out efficiently prevented the conversion of endogenous BV.

### 3D-PAULM of DrBphP expression in Blvra-/- brain

Since Blvra-/- significantly enhanced the performance of BphP-derived iLight in primary neurons, we next evaluated this Blvra-/- mouse model for deep brain imaging using our newly developed 3D-PAULM (**Fig. 2a**, see **Methods** for details). Briefly, 3D-PAULM utilizes a semispherical ultrasound transducer array with a central frequency of 4 MHz and can provide whole-brain imaging with the scalp and skull intact and a volumetric field of view of 8 mm in diameter (**Supplementary** Figs. 4a-b). 3D-PAULM is capable of two imaging modes: 3D RS-PAT and 3D ULM. The 3D RS-PAT mode repeatedly switches the BphP-derived molecular probes between the ON-state and OFF-state, and extracts the differential images of the probes while suppressing the background non-switching signals mostly from blood (**Fig. 2b, Supplementary** Fig. 4c). 3D RS-PAT has achienved a spatial resolution of ∼250 µm along the lateral dimension and ∼350 µm along the axial dimension (**Supplementary** Fig. 4d). The 3D ULM mode utilizes singular value decomposition as the spatial-temporal filter to extract the signals from the intravascularly injected gas-filled microbubbles^31^. By localizing the centroids of these microbubbles and accumulating over a large number of microbubbles (**Fig. 2b, Supplementary** Fig. 5), ULM can map detailed images of the blood vessels with sub-diffraction resolutions of ∼40 µm along all dimensions, about 10-fold improvement over the ultrasound Power Doppler image (**Figs. 2c-d**, **Supplementary** Fig. 6). The two imaging modes are seamlessly compatible and share the same 2D semispherical array, with complementary contrast mechanisms and comparable imaging depth.

**Figure 2.**
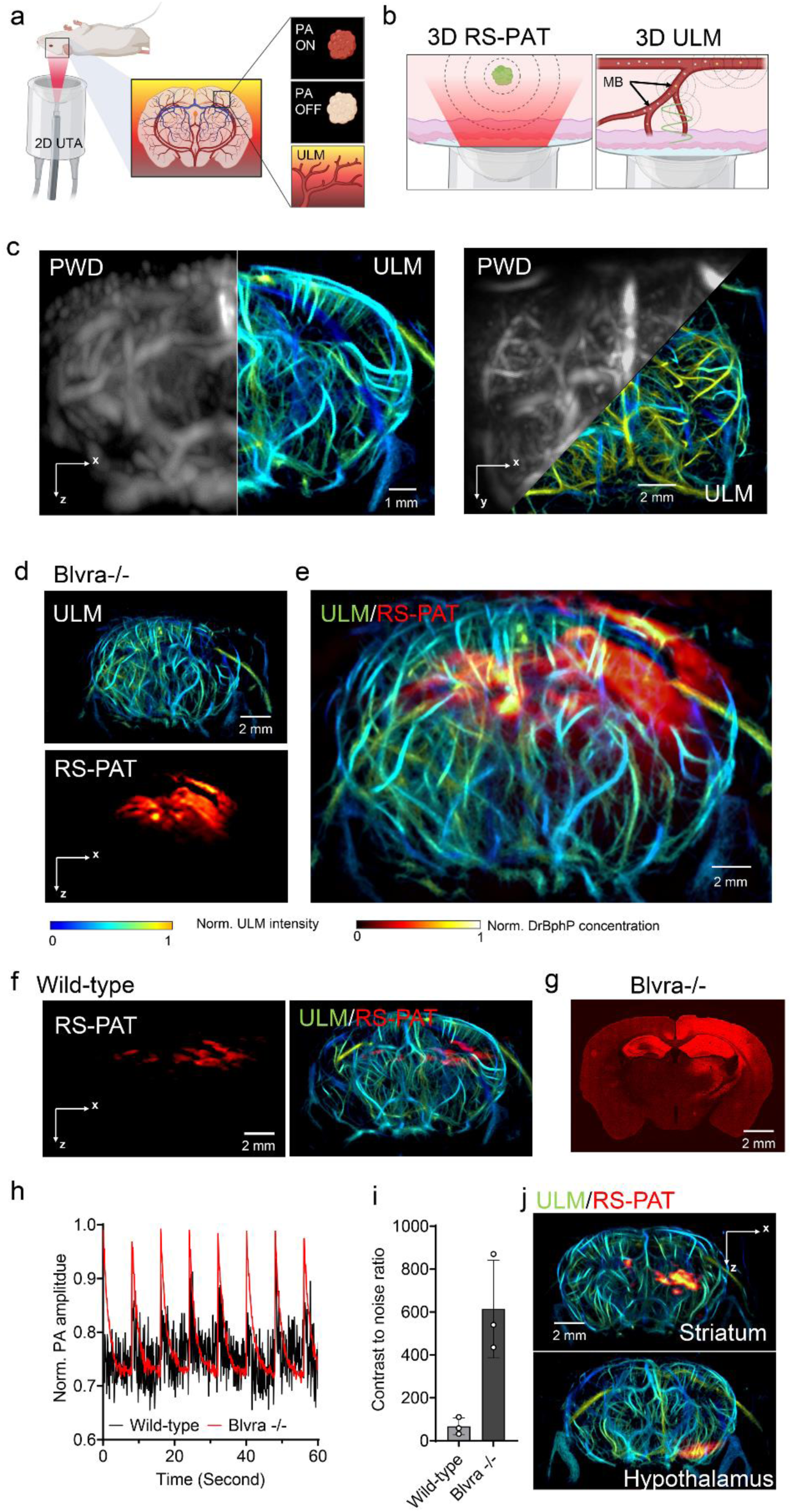
3D hybrid photoacoustic and ultrasound tomography (3D-PAULM) of neurons in the deep brain with skin and skull intact. **(a)** Schematic of the 3D-PAULM system with the 2D semispherical ultrasound transducer array (UTA). 3D-PAULM is capable of reversible-switching photoacoustic tomography (RS-PAT) and ultrasound localization microscopy (ULM). **(b)** Imaging principle of RS-PAT and ULM. RS-PAT detects the light-induced acoustic signals from the photoswitching BphP-based molecular probes. ULM localizes and tracks gas-filled microbubbles (MBs) in the blood stream. **(c)** Side-by-side comparison of the mouse brain vasculature imaged by ultrasound power Doppler (PWD) and ULM, projected along the coronal (*x-z*) or transverse (*x-y*) dimension. **(d)** 3D-PAULM of the DrBphP-PCM-expressing neurons in the hippocampus of the Blvra-/- mouse and the WT mouse. The ULM image shows the brain vasculature and the RS-PAT image shows the DrBphP-PCM-expressing neurons. (**e**) Overlay of the ULM and RS-PAT images. **(f)** 3D-PAULM of the DrBphP-PCM-expressing neurons in the hippocampus of the WT mouse. (**g**) Confocal microscopy image of the DrBphP-PCM expression in Blvra-/- brain slice. (**h**) Photoswitching dynamics of the RS-PAT signals in the Blvra-/- and WT mouse brains. **(i)** Contrast to noise ratio of the DrBphP-PCM signals in the Blvra-/- and WT brains. Data are presented as mean values +/-SD (*n*=3 independent injection sites). **(j)** 3D-PAULM images of the DrBphP-PCM-expressing neurons in the striatium and hypothalamus of the Blvra-/- mouse brain. We also extracted the mouse brain after the non-invasive PAT imaging experiment, and the isolated mouse brain (**Supplementary** Fig. 7b) shows the greenish color of phytochrome-expressing brain tissue.

We selected PCM of BphP from *D. radiodurans* (DrBphP-PCM: 55 kDa), which photoswitches between Pr (*λ*_max_= 660-698 nm; OFF state for PAT) and Pfr (*λ*_max_=740-780 nm; ON state for PAT) states^32^, as an imaging probe for 3D RS-PAT (**Supplementary** Fig. 7a). We targeted the DrBphP-PCM expression to the mouse hippocampus (depth from the skin surface: ∼2-3 mm) of the right hemisphere on both WT and Blvra-/- mice and performed 3D-PAULM on day 19 after AAV injection (**Supplementary** Fig. 7b). Here, the WT mice refer to animals having two intact copies of Blvra in genome (Blvra+/+), including transgenic mice available used in different experiments (see **Methods** for details). For the Blvra-/- mice, the 3D ULM image shows the detailed 3D vasculature of the brain, and the 3D RS-PAT image clearly delineates the AAV-mediated DrBphP-PCM expressing neurons of the right hippocampal region, with slight breach into the right cortex and left hippocampus (**Fig. 2e**). It is worth noting that the brain vasculature image by 3D ULM provided brain structural information such as the cortical diving vessels and hippocampus that are not accessible with 3D RS-PAT. Based on our data the PAT signals of DrBphP-PCM in the WT brain were substantially weaker than that in the Blvra-/- mouse (**Fig. 2f**). We compared the DrBphP-PCM one-photon fluorescence *in vivo* (ex: 665 nm; em: 750 nm) and observed ∼10-fold increase in the intensity of the emitted signal from the Blvra-/- mouse over the WT mouse (**Supplementary** Fig. 9a). The *ex-vivo* confocal microscopy of the mouse brain slices also confirmed the 3D RS-PAT result (**Fig. 2g**), showing the distribution of DrBphP-PCM-expressing neurons in both hemispheres highly consistent with the 3D RS-PAT results. The photoswitching efficiency of the Blvra-/- brain was ∼30%, whereas the WT brain had only ∼8% (**Fig. 2h; Supplementary** Fig. 8b). Note that the photoswitching efficiency is defined as the ratio of PAT signals at the ON- and OFF-state images. By taking the difference between the ON- and OFF-state images, the differential image can highlight the DrBphP-PCM signals while effectively removing the background hemoglobin signals (**Supplementary** Fig. 8a). Thus, the differential image of the DrBphP-PCM expressing neurons in the Blvra-/- brain showed a contrast to noise ratio of ∼615 at the right hippocampal region, which is about 9.2-fold of the WT mouse (**Fig. 2i**). To our best knowledge, this is the first time that PAT has been able to image genetically-labeled neurons in the deep brain with the skin and skull intact (∼3 mm deep from the scalp surface^33-35^). It is worth noting that our PAT is not able to resolve single neurons at such depths, due to the limited spatial resolutions. We were not able to quantitatively measure the DrBphP-PCM level in the mouse brains; however, based on our phantom data on the purified BphP proteins^11^, we estimated that the expression level was several micromoles.

Moreover, we targeted the DrBphP-PCM expression in neurons at even deeper brain regions, including the striatum (depth from the skin surface: ∼4-5 mm) and hypothalamic region (depth from the skin surface: ∼6-7 mm) (**Fig. 2j; Supplementary** Fig. 9b). From hippocampus to hypothalamus, 3D RS-PAT was able to clearly image the DrBphP-PCM-expressing neurons at the AAV injection regions, demonstrating the deep-brain imaging capability with the Blvra-/- mice (**Supplementary** Fig. 10). We would like to highlight that Blvra knockout has allowed for the high-sensitivity PAT of DrBphP-labelled neurons in deep mouse brains, which was not achieved before by using visible-light molecular probes, including GCaMPs.

### Blvra-/- increases the fluorescence level of miRFP720 and DrBphP in the brain

We next evaluated whether the fluorescence of BphP-derived NIR FPs, like miRFP720^8, 36^, is also increased in Blvra-/- mouse, with both one-photon and two-photon excitation. We compared miRFP720 fluorescence with one-photon excitation in primary neurons and live mice of WT and Blvra-/- genotypes. miRFP720 encoded under CAG promoter was delivered using AAV8. After 10 days of expression, Blvra-/- neurons exhibited a 2-fold increase in brightness (**Supplementary** Fig. 11a-d**; Supplementary** Fig. 12a). Similar enhancement of miRFP720 fluorescence was observed in skin fibroblasts of Blvra-/- mice (**Supplementary** Fig. 12b).

We next estimated how this improvement in brightness affected the penetration depth of *in vivo* fluorescence imaging of the brain. We first tested the AAV-mediated expression of miRFP720 by local injection in the dorsal hippocampus (depth from the skin surface: ∼2-3 mm) of the right hemisphere (**Supplementary** Fig. 11e). One-photon fluorescence (Ex. 665 nm; Em. 750 nm) of miRFP720 in the Blvra-/- mouse brains was ∼1.7-fold of that of WT mice, mainly in the right hemispheres (**Supplementary** Fig. 11f). We further tested the whole-brain expression of DrBphP-PCM using the AAV-PHP.eB capsid^37, 38^, which provides increased gene transfer to the CNS via vasculature (**Supplementary** Fig. 13a). After systemic delivery via tail vein injection, AAV-PHP.eB penetrates the blood-brain-barrier and transduces the neurons across the entire brain with enhanced expression per cell^38^. On day 17 after AAV8 transduction, the Blvra-/- mice displayed ∼2-fold higher one-photon fluorescence signals (Ex. 660 nm; Em. 750 nm) compared with the WT mice, with signals from the entire brains (**Supplementary** Fig. 13).

Finally, we investigated the feasibility of two-photon excited fluorescence imaging of miRFP720 expressed in the Blvra-/- mouse brain. The two-photon exciation of miRFP720 is peaked around 1280 nm in the NIR-II window, which can futher improve the imaging depth by reducing the light scattering and background autofluorescence. We injected the miRFP-720 AAV at 0.8 mm beneath the brain surface of the Blvra-/- mouse with a transcranicl window, and imaged the brain *in vivo* with two-photon microscopy (Ex. 1280 nm; Em. 720 nm) (**Fig. 3a**). The two-photon image of the miRFP720 expression clearly shows that a maximum imaging depth of 2.2 mm can be achieved in the Blvra-/- brain (**Fig. 3b**), as a result of strong miRFP720 signals and the deep NIR-II light penetration. The *ex vivo* image of the Blvra-/- brain confirmed the strong miRFP720 expression at depths. It is also worth noting that even at ∼2 mm depth, two-photon microscopy can achieve nearly single-cell resolution (**Fig. 3b**). By contrast, the two-photon fluorescence signals of the wild-type mouse were much weaker than the Blvra-/- mouse (**Fig. 3c**), with a maximal imaging depth of 1.3 mm. The depth-dependent averaged signal decay also demonstrates that the Blvra-/- mice can provide stronger two-photon fluorescence signals and thus larger imaging depths (**Fig. 3d**). Overall, the two-photon fluorescence singals of the Blvra-/- mice were ∼2.5-fold strongr than that of the WT mice (**Fig. 3e**, **Supplementary** Fig. 14).

**Figure 3.**
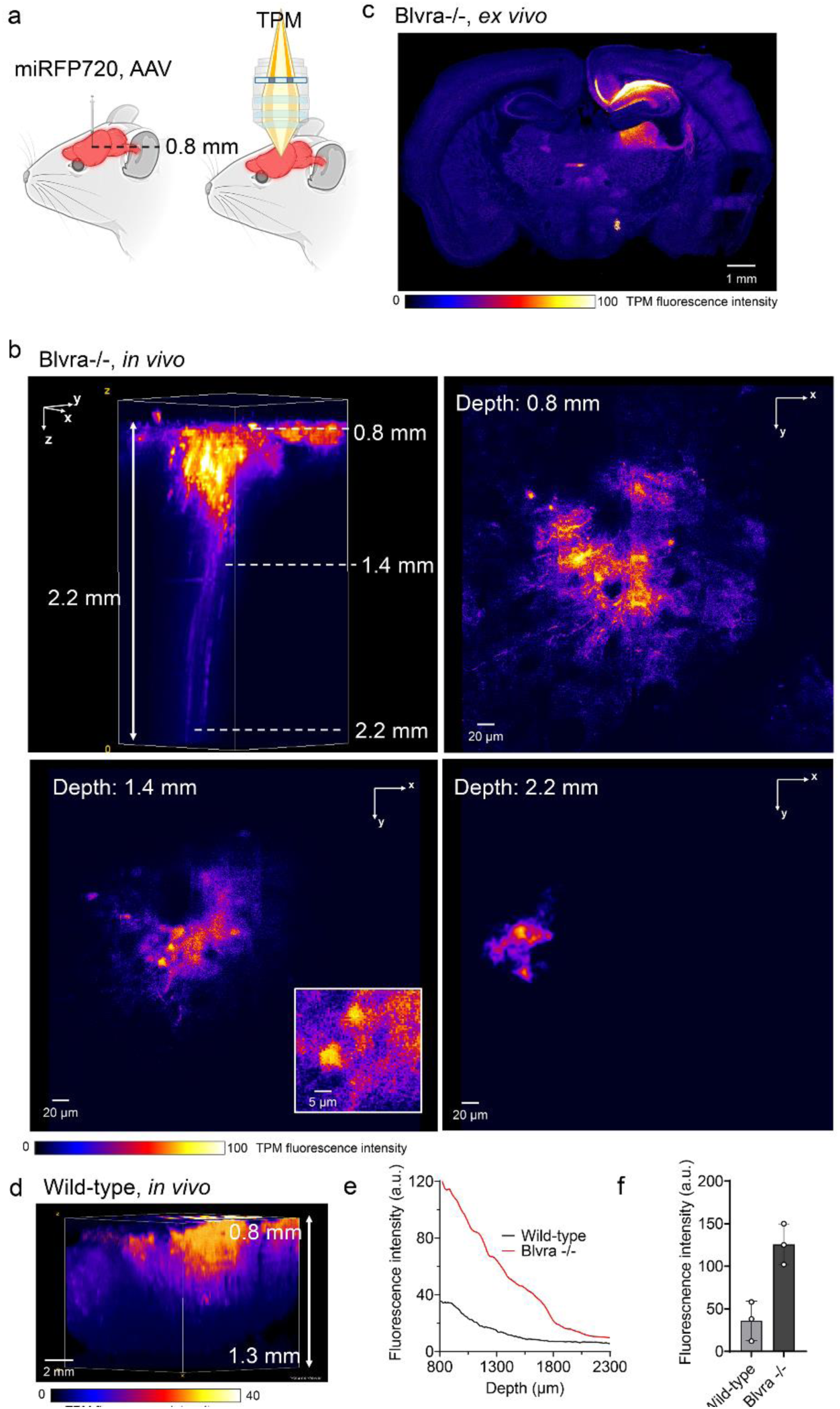
Two-photon microscopy of AAV-mediated miRFP720 in mouse brain. **(a)** Schematic of the AAV-mediated expression of miRFP720 injected at 0.8 mm depth beneath the brain surface. **(b)** Volumetric two-photon micrsocopy image of the miRFP720-expressing brain of the Blvra-/- mouse, showing a maximum imaging depth of ∼2.2 mm. Three representative images at 0.8 mm, 1.4 mm, and 2.2 mm depth are also presented. The inset image at the 1.2 mm depth showcases two individual cells. **(d)** Volumetric two-photon micrsocopy image of the miRFP720-expressing brain of the wild-type mouse, showing a maximum imaging depth of ∼1.3 mm. **(e)** Representative depth-dependent two-photon fluorescence signal decay for the Blvra-/- mouse and the wild-type mouse. **(f)** Depth-averaged two-photon fluorescence signals of miRFP720 of the Blvra-/- and wild-type mice (n = 3 independent injection sites).

### 3D-PAULM of AAV-Cre mediated BphP1 expression in liver and spleen

We next explored the whole-body deep-tissue imaging of the internal organs using a double-homozygous loxP-BphP1 Blvra-/- mouse. This mouse encodes loxP-EGFP-loxP-BphP1- mCherry-TetR construct and enables Cre-recombinase-mediated expression of BphP1-mCherry-TetR fusion protein in various tissues using the broadly available Cre driver mice or AAVs encoding Cre recombinase. We previously demonstrated RS-PAT of full-length *R. palustris* BphP1-expressing liver and spleen on loxP-BphP1 mice with relatively high contrast^14^. Here we imaged the AAV-Cre-mediated BphP1 expression in the liver and spleen of the loxP-BphP1 Blvra-/- mice to evaluate how Blvra knock-out affected the BphP1 detection by PAT in different tissues deep inside the mouse body. BphP1 fluorescence of the Blvra-/- liver was ∼2-fold stronger than that of the Blvra+/+ liver (**Fig. 4a**). The rich vasculature in the liver was clearly imaged by 3D ULM, and the BphP1-expressing hepatic cells were captured by the 3D RS-PAT with contrast to noise ratio of ∼440 at the center of the AAV injection region (**Fig. 4b**), which was ∼2.3-fold higher compared to Blvra+/+. The BphP1 expression was heterogeneous in the liver with multiple focal regions spread out of the injection site, which might reflect the liver zonation with varied metabolic levels^39, 40^. Similarly, the fluorescence of the Blvra-/- spleen was ∼2-fold higher compared to Blvra+/+ (**Fig. 4c**), and the 3D RS-PAT of the BphP1 signals had contrast to noise ratio of ∼504, which was ∼2-fold of the Blvra+/+ (**Fig. 4d**). The volumetric rendering of the BphP1 signals in the liver shows that the expression was more confined to the AAV injection site (**Fig. 4e**). The *ex vivo* fluorescence images of the extracted liver and spleen confirmed the RS-PAT results (**Supplementary** Fig. 15).

**Figure 4.**
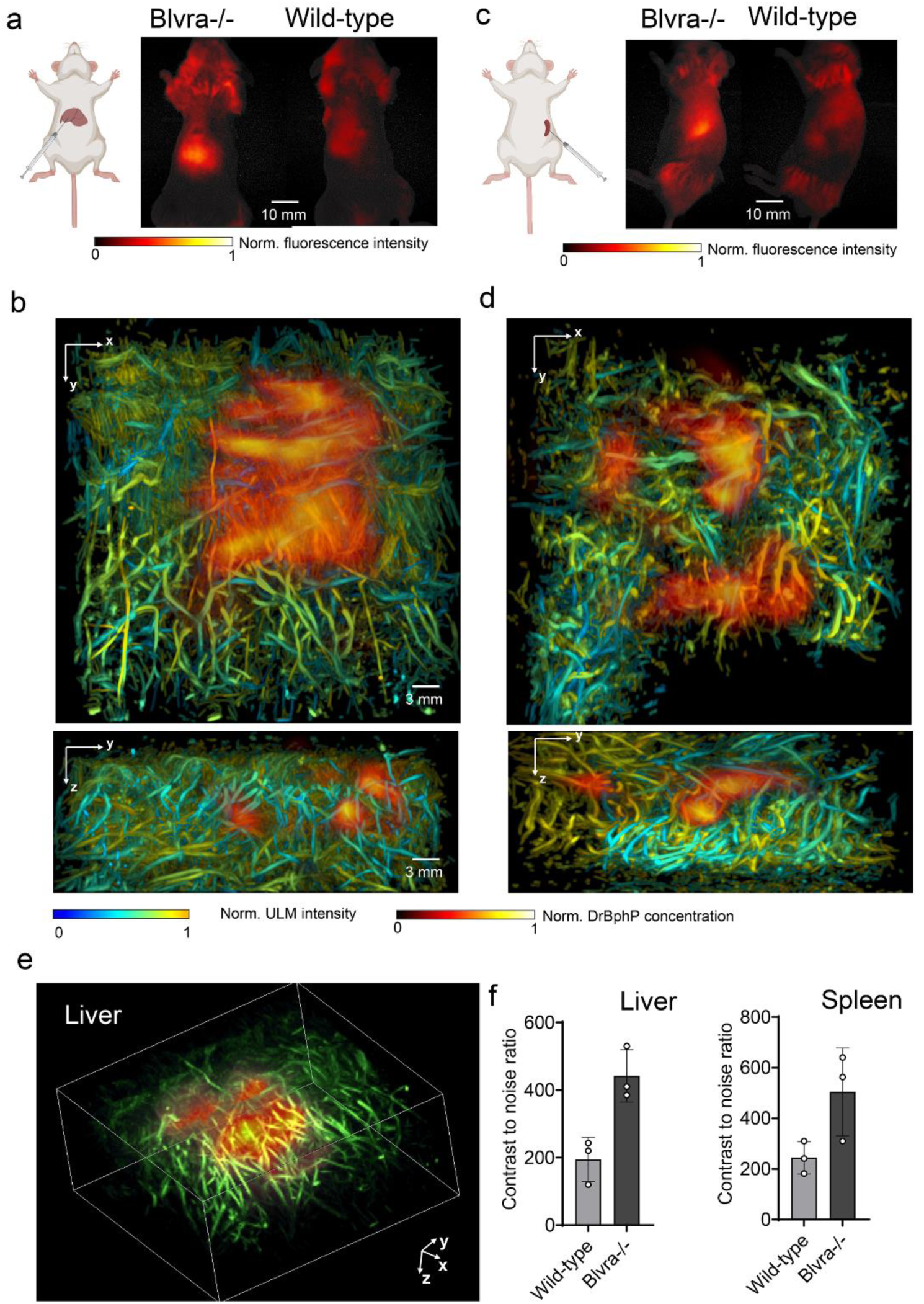
3D-PAULM of AAV-Cre mediated RpBphP1 expression in liver and spleen. **(a)** Schematic of Cre-mediated tissue-specific expression of BphP1 in *loxP*-BphP1 mice, and the wide-field fluorescence images of the AAV-Cre mediated RpBphP1 expression in the liver of Blvra-/- and Blvra+/+ mice. **(b)** 3D-PAULM images of the RpBphP1 expression in the liver of the Blvra-/- mouse. Two projections along the *x-y* and *y-z* planes are presented. **(c)** Wide-field fluorescence images of the RpBphP1 expression in the spleen of *loxP*-BphP1 Blvra-/- and *loxP*-Bphp1 Blvra+/+ mice. **(d)** 3D-PAULM projection images of the RpBphP1 expression in the spleen of the *loxP*-BphP1 Blvra-/- mouse. **(e)** 3D-PAULM volumetric rendering of the RpBphP1 expression in the liver of the *loxP*-BphP1 Blvra-/- mouse. (**f**) Contrast to noise ratio of RpBphP1 signals in the Blvra-/- and Blvra+/+ liver and spleen. Data are presented as mean values +/- SD (*n*=3 independent injection sites).

Thus, the Blvra-/- liver and spleen have a moderate improvement in 3D RS-PAT of BphP1 signals (**Fig. 4f**), when compared with that in the neurons. However, differences in BphP1 signals between organs were reduced in the Blvra-/- mice, which might be beneficial for studying the molecular activities at the whole-body level.

### 3D-PAULM of implanted transgenic tumors

It was shown that BV level in blood plasma is elevated in Blvra-/- mice^27^ and likely can reach the extravascular space and interstitial fluid. We next studied whether elevated BV from blood circulation could supply the implanted tumors, known to be highly vascularized with leaky vessels, and by this means enhance RS-PAT of BV-dependent imaging probe DrBphP-PCM expressed by the tumor cells (**Fig. 5a**). We allografted two types of 4T1 mouse breast cancer cells into the mouse mammary gland, the WT 4T1 and the DrBphP-PCM-expressing 4T1 (**Fig. 5b).** Both types of 4T1 cells have functional Blvra, but the performance of DrBphP can benefit from the extra BV supplied by the host circulation system of the Blvra-/- mice (**Fig. 5b**). We imaged the 4T1 tumors on the WT and Blvra-/- mice two weeks after implantation, and found that DrBphP-4T1 cells have stronger photoswitching performance in the Blvra-/- mice (**Fig. 5c**). The 3D ULM images can clearly outline both types of tumors based on the substantially elevated tumor vasculature density (i.e., angiogenesis) (**Fig. 5d**). 3D RS-PAT has better specificity on the DrBphP-4T1 tumors, whereas WT tumors are not visualized at all (**Fig. 5e**). By contrast, the DrBphP1-4T1 tumor in WT (Blvra+/+) mice had much weaker signals deep inside the tumor than the boundary region. Such difference in the signal distribution is also evident from the 3D RS-PAT results (**Fig. 5f**). The 3D RS-PAT results were confirmed by the fluorescence images *in vivo* and *ex vivo* (**Fig. 5g; Supplementary** Fig. 16). The contrast to noise ratio of the DrBphP-4T1 tumor in Blvra-/- mice was ∼3.6-fold higher than that in WT (Blvra+/+) mice (**Fig. 5h**), which is resultant from the elevated blood BV supply. Interestingly, confocal microscopy showed that the DrBphP1-4T1 tumor in the Blvra-/- mice had relatively uniform signals throughout the tumor, most likely due to the elevated interstitial BV level (**Supplementary** Fig. 16b).

**Figure 5.**
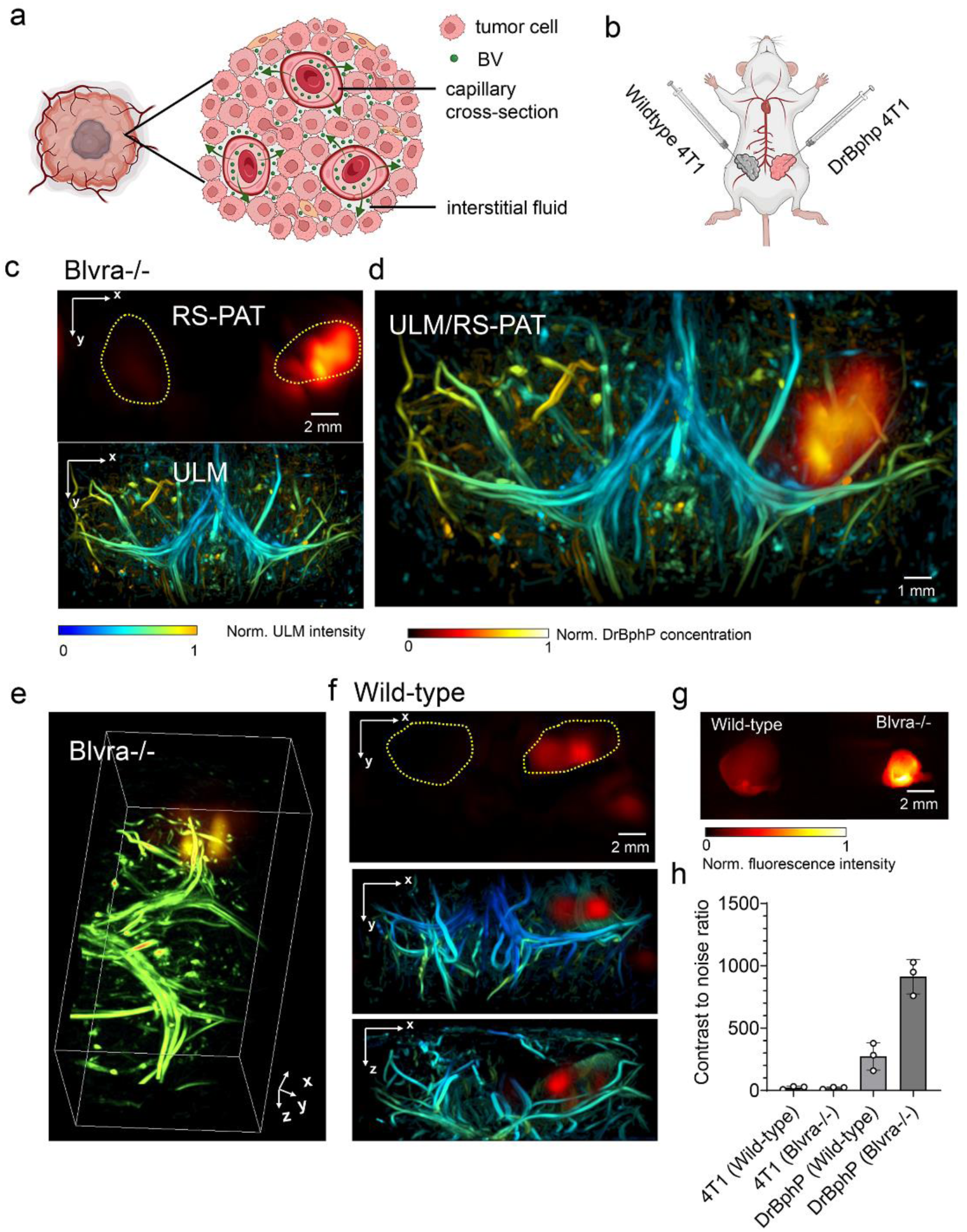
3D-PAULM of implanted transgenic tumors. **(a)** Illustration of the BV supply from blood circulation of Blvra-/- mouse inside the WT implanted tumor. **(b)** Illustration of implanting DrBphP-PCM-expressing 4T1 and WT 4T1 tumors in the Blvra-/- and WT mice. **(c)** 3D-PAULM projection images of the implanted 4T1 tumors in the Blvra-/- mouse. The yellow dashed lines outline the tumor regions based on the elevated vascular density in the ULM image. Only the DrBphP-PCM-4T1 tumor is visible in the RS-PAT image. Note that the DrBphP-PCM signal levels are relatively uniform throughout the tumor, as indicated by the double yellow arrowheads. (**d**) Overlay of the RS-PAT and ULM images. **(e)** 3D-PAULM volumetric rendering of the DrBphP-PCM-expressing 4T1 tumor in the Blvra-/- mouse. **(f)** 3D-PAULM projection images of the implanted 4T1 tumors in the WT mouse. Note that the DrBphP-PCM-4T1 tumor signals are much weaker than those in the Blvra-/- mouse. Moreover, the DrBphP-PCM signals are stronger on the boundaries, as indicated by the double yellow arrowheads. **(g)** Wide-field fluorescence images of the DrBphP-PCM-expressing 4T1 tumors extracted from the Blvra-/- and WT mice. **(h)** Contrast to noise ratio of 4T1 tumors in the Blvra-/- and WT mice. Data are presented as mean values +/- SD (*n*=3 independent mice).

### Light-mediated control of insulin production in type 1 diabetes

We next studied the NIR optogenetic transcription activation for correction of pathological conditions, such as type 1 diabetes (T1D), when the production of insulin is abolished by loss of pancreatic β cells. We first validated the mouse insulin production in cultured HeLa cells (**Fig. 6a-c**). Cells were co-transfected with pCMVd1-NLS-Gal4-iLight-VP16 and pG12-Proinsulin plasmids in a 1:3 ratio, supplemented with BV, and irradiated with 660 nm light or kept in darkness. Cell medium collected after 48 h was used to determine secreted mouse insulin by ELISA. Medium from illuminated cells contained 3.1 µg/l insulin, whereas cells kept in darkness did not produce detectable insulin levels (ELISA’s detection limit is 0.2 µg/l insulin) (**Fig. 6c**). We used 10 times concentrated sample of culture medium collected from HeLa cells, in line with unconcentrated samples to verify that we have confirmed insulin production, since insulin produced by the monolayer of HeLa cells was quite low for detection.

**Figure 6.**
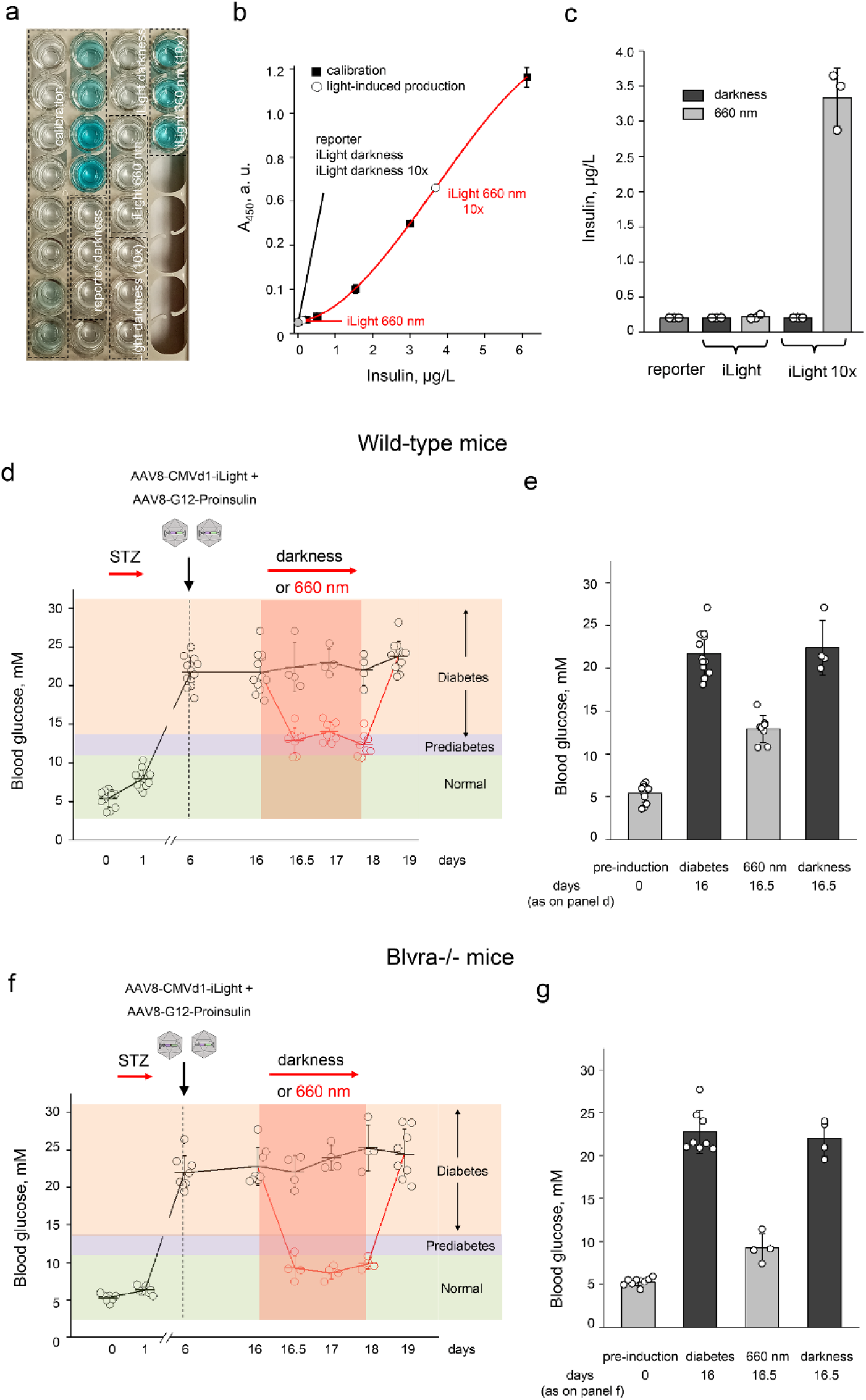
Optogenetic therapy of type 1 diabetes (T1D) using light-induced insulin production. **(a-c)** Testing the iLight-mediated insulin production in HeLa cells. For light-induced insulin production, HeLa cells were co-transfected with pCMVd1-NLS-Gal4-iLight-VP16 plasmid (or empty pcDNA3.1+ plasmid) and reporter pAAV2-G12-Proinsulin plasmid in 1:4 ratio. Cells were illuminated with a 660/15 nm LED array (0.4 mW cm^-2^) or kept in darkness for 48 h. Insulin concentration in cultured media was determined using a mouse insulin ELISA kit with a colorimetric readout. The ELISA plate with samples of culture medium collected from transfected HeLa cells after 48 h to determine secreted insulin (a), calibration curve (b), and detected insulin levels for various conditions (c) are shown. **(d-f)** Glucose dynamics in T1D mice during the development of the pathology and following the optogenetic manipulation with NIR light. T1D was induced by repeated injection of streptozotocin (STZ, 40 µg/kg of body weight for 3 days). After confirmation of diabetes (see the colored concentration ranges), mice were co-injected with AAV8 encoding CMVd1-NLS-Gal4-iLight-VP16 and/or G12-Proinsulin. After 10 days of transduction, mice were illuminated with a 660/15 nm LED array (3.5 mW cm^-2^) for 48 h to induce insulin production in the liver or kept in darkness. Fasting blood glucose levels in WT and Blvra-/- mice in the norm, before and after induction of T1D, and after optogenetic manipulation are shown in (f), ns, non-significant. Data were analyzed using one-way ANOVA and are presented as mean values +/- SD (n=4 WT mice kept in darkness, n=8 illuminated WT mice; n=4 Blvra-/- mice kept in darkness, n=4 illuminated Blvra-/- mice).

T1D in mice was induced by repeated injection of streptozotocin (STZ), a glucosamine-nitrosourea compound and alkylating agent produced by *S. achromogenes,* which damages DNA and is particularly toxic to islet β cells. We monitored the blood glucose level during STZ injections and used the STZ dosage of 40 µg/kg of body weight for 3 sequential days (**Fig. 6d,e**). This ensured the high animal survival rate with the final blood glucose level of 21.75 mM in WT and 22.79 mM in Blvra-/- mice (**Fig. 6f**). After confirming the T1D state, the WT and Blvra-/- mice were systemically co-injected with AAV8 encoding CMVd1-NLS-Gal4-iLight-VP16 and G12-Proinsulin. Systemically injected AAV8 transduces liver in mice^41^ and humans^42, 43^ with high efficiency. Following 10 days of expression, the mice were illuminated to enable liver insulin production or kept in darkness for 48 h (days 16-18 in **Fig. 6d,e**). During this period, blood glucose was monitored at 12 h, 24 h and 48 h. A clear trend of the dependence of fasting blood glucose level on the optogenetic manipulation was observed. iLight-mediated insulin production in the liver decreased the blood glucose in both WT and Blvra-/- mice, with lower blood glucose levels in Blvra-/- mice (12.93 mM in WT mice versus 9.26 mM in Blvra-/- mice, p<0.005, one-way ANOVA) (**Fig. 6f**).

A prediabetic state in mice is characterized by fasting blood glucose values between 11.1-13.8 mM ^44^. Our data evidence that optogenetically produced insulin improved the diabetic state (fasting blood glucose >13.8 mM) towards the borderline between diabetic/prediabetic state in WT mice but further decreased it to the normal blood glucose range of <11.1 mM in Blvra-/- mice.

## DISCUSSION

We found that the performance of the BV-dependent OT, fluorescent and PAT imaging probes, which all are derived from three different BphPs (DrBphP, RpBphP1 and IsPadC), is significantly enhanced in Blvra-/- mouse.

iLight-based optogenetic gene transcription enhanced 4-25 times in various primary cell types isolated from Blvra-/- mice over that in cells from WT mice, with, importantly, a 100-fold activation contrast in primary Blvra-/- neurons. Notably, exogenous BV did not improve the performance of iLight in Blvra-/- cells, suggesting that all OT molecules are assembled with endogenous BV, i.e. are in the holoform.

Optogenetic therapy of pathological conditions, such as T1D, using iLight-mediated insulin production decreased blood glucose to the normal range in Blvra-/- mice but not WT mice, thus managing the diabetic conditions non-invasively with NIR light. Not shown but worth mentioning that other pathological conditions related to an inability to produce metabolic proteins or hormones could be corrected similarly.

We developed a hybrid imaging system 3D-PAULM, which allowed both RS-PAT of the BphP-derived NIR molecular probes and high-resolution ULM of the vasculature deep inside the tissues. Although PAT can also image the blood vasculature by using hemoglobin as the endogenous contrast, it is clear that the vasculature image quality by ULM is superior to that by PAT, with 10-fold better resolution. This is mostly because ULM does not suffer from the limited-view problem as in PAT^45, 46^, and the ULM inherently has better spatial resolutions than PAT using the same transducer array. Therefore, we think that it is beneficial to utilize ULM to provide a high-quality vasculature image of the deep tissues.

Using 3D-PAULM, we imaged DrBphP-PCM-expressing neurons in Blvra-/- brain with a signal-to-noise ratio of ∼615, which is almost an order of magnitude higher than in WT mice. To our knowledge, this is the first time that PAT was able to image genetically-labeled neurons with such high contrast in the deep brain with the skin and skull intact. Moreover, we were able to look deeper into the brain of Blvra-/- mouse, clearly imaging DrBphP-PCM-expressing neurons in the striatum (4-5 mm depth) and hypothalamic region (6-7 mm depth).

Using 3D-PAULM, we imaged Cre-mediated expression of genetically-encoded RpBphP1 in the liver and spleen and found that even tissues naturally enriched with BV chromophore exhibit enhanced imaging performance in Blvra-/- animals. WT breast tumors allografted into Blvra-/- mouse were enriched with the biliverdin via blood supply and exhibited a contrast-to-noise ratio of ∼3.6-fold over the WT mice, making the Blvra-/- mouse a beneficial model for non-invasive studies of tumorigenesis and tissue transplants using various BphP1-derived NIR imaging modalities.

Unlike DrBphP and BphP1 used in our 3D-PAULM, some BV-dependent imaging probes evolved toward higher binding efficiency of BV chromophore. This is the case of miRFP720 NIR FP, which was derived from *R. palustris* BphP and subjected to directed molecular evolution^8, 36^. We have shown that despite the substantial optimization, miRFP720 still exhibited twice higher fluorescence in the Blvra knock-out brain tissues enriched with BV.

Our results have also shown that Blvra-/- mice can provide much brighter fluorescence for both miRFP720 and DrBphP, when compared with the WT mice. This is true for both one-photon fluorescence with the excitation wavelength around 665 nm and two-photon fluorescence with the excitation around 1280 nm. Particularly, the two-photon microscopy has showcased an imaging depth of ∼2.2 mm, with nearly single-cell resolution. Attempts have been made earlier to improve the brightness of BV-dependent probes *in vivo* using systemic BV administration. Intravenous injection of 250 nmole (∼7 mg/kg) BV increased near-infrared fluorescent protein IFP1.1 fluorescence in the liver by ∼5-fold and did not display observable toxicity in mice^47^. However, this approach suffers from major limitations, such as fluctuating levels of BV and its uneven distribution in the target tissue, which makes quantitative studies doubtful; inability to perform longitudinal imaging or optogenetic manipulation without multiple BV administration; and poor penetration of BV through the blood-brain barrier, which makes brain unaccessible. Another way was to overexpress heme oxygenase and interference RNA for BV reductase in specific tissues. For example, it was shown that injection of AAVs with these both components resulted in the normal functioning of BphP-based OT in the mouse brain^48^. However, silencing with shRNA does not abolish all activity and the remaining protein, especially enzymes, can cope with the metabolic load.

BV formation primarily occurs in the mononuclear phagocyte system of the liver and spleen, which are responsible for the deposition of senescent red blood cells. Erythrocytes can also release hemoglobin after hemolysis due to autoimmune diseases, infections, or medications. In addition, in all cell types heme is available as a prosthetic group of hemoproteins. In muscle cells, myoglobin is a considerable source of heme, which can be released and sequestered in macrophages after tissue damage^49^. BV serves as an antioxidant and protects tissues by scavenging peroxyl radicals. In Blvra-/- mice the accumulation of BV and a notable decrease of bilirubin in blood plasma were reported^27^.

Blvra knockout mice are now widely available and used for studies of various aspects of heme metabolism. They are independently developed by several research groups^26-29, 50^, and also available from depositories of transgenic mice, including Jackson Laboratories (stock <u>036908)</u> and Japan National Institute of Biomedical Innovation (resource <u>nbio370</u>).

Blvra gene homozygous and heterozygous mutation have been documented in humans^51,52^. Heterozygous mutation in Blvra gene can cause hyperbiliverdinaemia when combined with decompensated liver cirrhosis. Individual with heterozygous mutation in Blvra gene had two children heterozygous for the identical mutation in the Blvra gene, with no clinical signs of liver disease and had normal levels of biliverdin^51^, likely indicating the sufficient compensatory role of one copy of functional Blvra. As was revealed in the first report of a homozygous Blvra inactivating mutation the complete absence of Blvra activity is a non-lethal condition^52^, the most evident phenotypic characteristic of which is the appearance of hyperbiliverdinaemia (green jaundice) accompanying cholestasis episodes. Moreover, many studies indicate a positive effect of the inactivation of Blvra to treat heme-metabolism-related human diseases. Neonatal jaundice caused by either reduced bilirubin conjugation or bilirubin overproduction) is observed in about 15% premature infants in low-/middle-income countries^53, 54^, results in acute bilirubin encephalopathy (kernicterus), cerebral palsy and, eventually, if untreated, death. In adults, a similar pathologic condition is an unconjugated hyperbilirubinemia, causing various neurologic dysfunctions. Currently, the only available treatments are intensive phototherapy and exchange transfusion. Inhibition of biliverdin reductase is considered to be a novel promising and safe treatment for unconjugated hyperbilirubinemia^29^. The beneficial anti-inflammatory, anti-apoptotic, anti-proliferative, and antioxidant properties of the elevated biliverdin levels have been documented in many disease models, including ischemia-reperfusion injury, organ transplantation, and immune responses^55^.

Built upon our proof-of-principal results, future work should include disease-driven studies based on this deep-tissue imaging platform, such as monitoring the neuronal degeneration in the ischemic storke and liver regeneration in the liver cancer therapy^56, 57^. Moreover, BphP-based NIR biosensors are greatly needed to empower the imaging technology for probing the cellular and molecular functions in deep tissues, including calcium-sensitive indicators for monitoring neuronal activities^58^, and surface-markers-specific reporters for targeting metastatic and dormant tumor cells^59^. Besides, 3D-PAULM can be futher improved by incorporating more super-resolution imaging strategies to enchance the image quality, including focused light delivery using the BphP-based probes as the wavefront-shaping guide stars^60^.

## METHODS

### Ethical statement

All procedures with animals were performed in an AAALAC-approved facility and received ethical approval from the Institutional Animal Care and Use Committee of the Albert Einstein College of Medicine (protocol 00001050) and Duke University (protocol A203-22-12).

### Housing and husbandry

Mice were maintained on a 12 h light and dark cycle at room temperature and standard humidity (∼40-55%), with ad libitum access to food and water. Same-sex littermates were housed together in cages with chopped corn cob bedding (2-5 mice per cage). Environmental enrichment included pieces of compressed cotton nestlets and paper huts.

### Molecular cloning

To make a pAAV-G12-AkaLuc plasmid, the fragment encoding AkaLuc was amplified from pAAV-TRE-Tight2-AkaLuc (Addgene #186186) and swapped with CheRiff-T2A-mCherry in pAAV-G12-CheRiff-T2A-mCherry (Addgene #170275) using BamHI and EcoRI restriction sites. Previously published proinsulin sequence^32^ was used to synthesize Proinsulin cDNA by GenScript. To make pAAV-G12-Proinsulin, it was digested with BamHI and BglII restriction enzymes and then ligated with pAAV-G12 backbone.

### Genotyping of Blvra-/- knock-out mice

Several strains of Blvra-/- mice were recently reported^26-29, 50^. Founders of Blvra-/- strain characterized here were a gift of Drs. Kenta Terai and Michiyuki Matsuda from Kyoto University, Japan and allowed us to establish our own colony. Blvra-/- mice were developed by a CRISPR/Cas9 system targeting the murine *Blvra* gene (NM_026678.4) as described earlier^28^. F0 mouse with large DNA-fragment inversion between the two gRNA-target sites was selected for further studies. F0 mouse with large DNA-fragment inversion between the two gRNA-target sites (Supplementary Fig. 2a) was selected and further crossed with B6 albino mice to obtain Blvra-/- progeny in F2 generation. Animals were obtained via JCRB Laboratory Animal Resource Bank of the National Institutes of Biomedical Innovation, Health and Nutrition (Tokyo, Japan), animal ID nbio-370. The presence of the transgene in the offspring was detected using PCR. Ear punches (∼1 mm in diameter) were incubated with 250 μl of 10 mM NaOH, 0.1 mM EDTA at 95°C for 10 min. One microliter or solution was used as a template for PCR. The thermal protocol consisted of an initial hold of 95°C for 1 min, followed by 35 cycles of 95°C for 15 s, 58°C for 20 s, 72°C for 30 s, and an additional 72°C for 2 min. The reaction volume was 20 μl. PCR products were visualized using agarose gel electrophoresis. Genotypes were determined using primers annealing to WT allele (5’-TGGTAGTGGTTGGTGTTGGC-3’ and 5’- CCACTACTCGGCATGGTTCT-3’, amplicon size 216 bp) and to the inverted allele (5’- CCACTACTCGGCATGGTTCT-3’ and 5’-AAAGGATGAAAGGCAACATGAGCAG-3’, amplicon size 417 bp) (**Supplementary** Fig. 2b). Since we established a colony of *loxP*-BphP1 homo, Blvra-/- mice we determined the progeny homozygous for BphP1 insert using primers annealing to intact *ROSA26* locus (5’-AGCACTTGCTCTCCCAAAGTC, 5’- TGCTTACATAGTCTAACTCGCGAC, product size 564 bp) and transgene floxed *EGFP* sequence (5’-GGCAGAGGATCGTTTCGCGG, 5’-GAAGCACTGCACGCCGTAGG, product size 240 bp) as in **Supplementary** Fig. 3b.

### Preparation of high-titer AAVs

AAV8 particles encoding G12-AkaLuc, G12-AkaLuc-P2A- Proinsulin, CMVd1-NLS-Gal4-iLight-VP16 or CAG-miRFP720 were prepared as described earlier^61^. Briefly, plasmids for AAV production were purified with NucleoBond Xtra Maxi EF kit (Macherey-Nagel) and AAV-293T cells (Agilent) were co-transfected with the pAAV2-G12- AkaLuc, pAAV2-G12-AkaLuc-P2A-Proinsulin, pAAV2-CMVd1-NLS-Gal4-iLight-VP16 or pAAV2-CAG-miRFP720, a pAAV2/8 Rep/Cap plasmid (Addgene#112864) and a pHelper plasmid using polyethyleneimine (Santa Cruz). Cell growth medium was collected 72 h after transfection. 120 h after transfection cells and growth medium were collected and combined with medium collected at 72 h. Cells were pelleted by centrifugation and then lysed with salt-active nuclease (SRE0015, Sigma-Aldrich). 8% PEG was added to the supernatant, incubated for 2 h on ice and then pelleted by centrifugation. PEG pellet was treated with SAN, combined with lysed cells and clarified by centrifugation. The supernatant was applied on an iodixanol gradient and subjected to ultracentrifugation for 2 h 25 min at 350,000 g. Virus fraction was collected, washed and concentrated on Amicon-15 100,000 MWCO centrifuge devices (Millipore). Purified virus aliquots were stored at -80°C. Virus titer was defined by qPCR. For this, virus aliquots were consequently treated with micrococcal nuclease (New England Biolabs) and proteinase K (Fermentas) and then used as a template for qPCR.

AAV2 and AAV8 encoding human Cre recombinase were purchased from the AAV Gene Transfer and Cell Therapy Cora Facility of the University of Helsinki (Finland). AAV8 encoding Cre recombinase and AAV9 encoding CAMKII-DrBphP-PCM were prepared by the Duke University Viral Vector Core in the Center for Advanced Genomic Technologies. These viruses were produced in accordance with previously published methods^61, 62^.

### Isolation of primary cultures

Primary cultures of neurons, endothelial cells, and fibroblasts were isolated from postnatal (P0-P1) *Blvra-/-* mice (both males and females). Cortical neurons were isolated using the published protocol^63^ and cultured in Neurobasal Plus Medium with B-27 Plus Supplement (Gibco), additional 1 mM GlutaMAX (Gibco, 35050061), 100 U/ml penicillin and 100 μg/ml streptomycin, on poly-D-lysine (EMD Millipore) coated glass coverslips (thickness 0.13-0.17 mm, diameter 25 mm, ThermoFisher Scientific) at density ∼70,000 cells per coverslip. Half of the medium was exchanged twice a week.

Primary fibroblasts were isolated from the skin using a protocol described earlier^64^, with the following modifications. Digestion was performed in DMEM with 10% FBS, 100 U/ml penicillin and 100 μg/ml streptomycin, 0.25% collagenase type I, 15 mM HEPES pH7.4 at 37°C for 1.5 h while shaking at 220 rpm. Cells were cultured in DMEM with 10% FBS, 100 U/ml penicillin and 100 μg/ml streptomycin. Upon reaching confluency the fibroblasts were dissociated with a trypsin-EDTA solution and replated at a 1:5 ratio. Under these conditions, the fibroblasts might be cultured for up to 6 passages (∼20 days).

Endothelial cells were isolated from the mouse lungs of neonatal mice. Lungs were rinsed twice in cold HBSS, microdissected into small pieces (∼0.5-1 mm diameter) and dissociated with collagenase type I (%) in high-glucose DMEM, 10% FBS, 100 U/ml penicillin and 100 μg/ml streptomycin, 15 mM HEPES pH7.4 at 37°C for 1 h with gentle rotation. After that suspension was passed through a 70 μM cell strainer, sedimented at 600 g for 5 min and washed once with high-glucose DMEM, 10% FBS, 100 U/ml penicillin and 100 μg/ml streptomycin and resuspended in high-glucose DMEM, 5% FBS,100 U/ml penicillin and 100 μg/ml streptomycin and cultured on the collagen I–coated (Gibco, A10483-01) 6-well plates. Cells were allowed to reach 70% confluency before transduction.

### AAV transduction of primary cells

On the 5th day *in vitro* (5 DIV) cells were transduced with two AAV8 delivering iLight^25^ and AkaLuc luciferase reporter gene^30^. As some leakage of the G12-AkaLuc construct may occur, we separately estimated the bioluminescence signal in cells transduced with AAVs encoding G12-AkaLuc and negative control iGECInano (AAVeq, **Fig. 1b**) at a similar ratio of 1:3 with a multiplicity of infection of 10^5^. This background signal was subtracted from the bioluminescence radiance evoked in cells transduced with full-component OT (iLight with AkaLuc, **Fig. 1b**). Immediately after transduction, where necessary, cells were supplemented with BV: the final concentration was 2 µM for neuronal cultures and 5 µM for all other cell types. Plates kept in darkness were protected with aluminum foil after transduction, and all subsequent manipulations were performed under deemed green light, which prevented iLight transition into the Pfr state.

### Hydrodynamic transfection of the liver

Nine WT and nine Blvra-/- mice (3-10 months of age, females) were used in the studies (n=3 for each of the 3 independent experiments). For hydrodynamic transfection of the liver, mice in the cage were pre-warmed at 37.5°C for 15 min using a heating pad to promote vasodilation. Mouse was restrained in Tailveiner Restrainer from Braintree Scientific (Braintree, MA, USA) and 16.6 μg of pCMVd1-NLS-Gal4-iLight-VP16 and (or empty pcDNA3.1+ plasmid) with 43.4 μg of pAAV2-G12-AkaLuc reporter plasmid in 1-2 mL warm DPBS (equivalent to 7% of mouse body weight) were injected through the tail vein within 5-7 s. For mice kept in darkness all subsequent manipulations were performed under a 520 nm safelight. Mice were allowed to recover for 1 h and then were illuminated with a 660/15 nm LED array at 3.5 mW cm^-2^ or kept in darkness for 24 h (**Fig. 1f**). Bioluminescence imaging was performed using IVIS Spectrum after injection of AkaLumine-HCl substrate (75 nmol/g of body weight).

### In vivo fluorescence and bioluminescence imaging

Fluorescence and bioluminescence imaging was performed on the IVIS Spectrum instrument (PerkinElmer). Throughout the *in vivo* imaging session animals were maintained under anesthesia with 1.5% vaporized isoflurane. The instrument stage was temperature-controlled (37°C) to avoid hypothermia. For EGFP imaging 465/20 nm excitation and 520/30 nm emission filters were used. For AkaLuc bioluminescence imaging *in vivo,* the animals were injected with NIR-emitting luciferin analog AkaLumine-HCl (Sigma-Aldrich, 808350, 0.075 nmol/g of body weight) 15 min before imaging. For bioluminescence detection *in vitro* primary cells were supplemented with 250 μM AkaLumine-HCl. Detection was performed in bioluminescence mode with an open emission filter. Data were analyzed using Living Image v.4.3 software (Perkin Elmer/Caliper Life Sciences) and plotting and statistical analysis were performed using OriginPro v.9.7.188.

### Type 1 diabetes model

WT and Blvra-/- mice (8- to 10-week-old), were fasted for 16 h and injected with streptozotocin (STZ, Sigma, S0130), 40 µg/kg of body weight for 3 sequential days. STZ was diluted in citrate buffer (pH 4.28) and injected intraperitoneally within 1 min after dilution. First two days after STZ injection 10% sucrose water was provided to prevent possible episodes of hypoglycemia. Fasting blood glucose samples were taken before mice were released from the illumination cage for feeding (12 h fasting, see description for illumination experiment). Glucose level was measured using Contour Next EZ Glucometer.

### Optogenetic transcriptional activation with NIR light

Ten days after viral transduction cells were illuminated with 660/15 nm light (0.4 mW cm^-2^) or kept in darkness for 48 h (**Fig. 1a**). For bioluminescence imaging, cells were detached using trypsin (neurons) or collagenase I (fibroblasts, endothelial cells) and washed once with PBS. Cells were resuspended in PBS with 5% FBS, transferred into the black 96-well plate and supplemented with 250 µM AkaLumine-HCl (TokeOni, Sigma, 808350), a luciferin analog displaying bioluminescence peak at 677 nm. Detection was performed on the IVIS Spectrum instrument (PerkinElmer/Caliper Life Sciences) in bioluminescence mode with an open emission filter. AkaLuc expression was estimated based on the average bioluminescence radiance of the circular ROIs corresponding to the perimeter of the wells.

For light-induced insulin production, HeLa cells were cultured in DMEM with 10% FBS, 100 U/ml penicillin and 100 μg/ml streptomycin. Cells were transfected with 0.08 µg of pCMVd1-NLS-Gal4-iLight-VP16 (or empty pcDNA3.1+ plasmid) and 0.32 µg of pAAV2-G12-AkaLuc (0.4 µg per well) using Effectene transfection reagent (Qiagen). Culture media was changed 8 h after transfection. Cells were illuminated with 660/15 nm light (0.4 mW cm^-2^) or kept in darkness for 48 h. Insulin level in cultured media samples was detected using a commercial mouse insulin ELISA kit (Mercodia, 10-1247-01).

For *in vivo* experiments, mice were placed in conventional transparent cages without bedding, and illuminated from the bottom with a 660/15 nm LED array (**Fig. 1f**) with the intensity of the activation light 3.5 mW cm^-2^. For better illumination and imaging, the belly fur was shaved with a trimmer. Control animals were kept in the same cages in complete darkness. Every 12 h mice were transferred for rest in conventional cages with free access to food and water (1 h).

### Epifluorescence microscopy

For fluorescence microscopy neurons were plated on poly-D-lysine precoated MatTek glass bottom dishes (35 mm). Cells were transduced with AAV8 CAG- miRFP720 (multiplicity of infection of 10^5^) on 5 DIV and cultured in Neurobasal Plus Medium with B-27 Plus Supplement (Gibco), and additional 1 mM GlutaMAX (Gibco, 35050061), 100 U/ml penicillin and 100 μg/ml streptomycin for 12 days before imaging. Where necessary, cells were supplemented with 2 µM BV 24 h before imaging. The fluorescence of miRFP720 was recorded in the Cy5.5 channel for WT and Blvra-/- cells using the same acquisition settings. The Olympus IX81 microscope was operated with a SlideBook v.6.0.8 software (Intelligent Imaging Innovations).

### Flow cytometry

Flow cytometry analysis of primary cortical neurons and skin fibroblasts was carried out on Cytek Aurora (neurons) and BD LSRII (skin fibroblasts) flow cytometer. Before analysis, cells were gently detached using trypsin, and suspended in cold PBS, 4% FBS with 2 mM EDTA. At least 50,000 cells per sample were recorded. The fluorescence intensity of miRFP720 expressing cells was analyzed using the AlexaFluor 700 channel. The data were analyzed using FlowJo v.7.6.2 software.

### Hybrid 3D photoacoustic and ultrasound localization microscopy system (3D-PAULM)

The 3D-PAULM system comprises a programable ultrasound scanner (Vantage 256, Verasonics, Kirkland, WA), multiple pulsed lasers for PAT excitation, and a customized 2D semispherical ultrasound transducer array (Imasonics, France) for both PAT and US imaging. The semispherical transducer array has 256 elements, with a central opening for inserting an optical fiber bundle. The transducer array has a center frequency of 4 MHz, with a −6 dB bandwidth of ∼45% in the transmit/receive mode, although it was fine-tuned to maximize the receiving bandwidth. The transducer array has a curvature radius of 40 mm and a total aperture size of 57 mm. The well-sampled 3D field of view (3D-FOV) of the transducer (or the focal zone) is 8×8×8 mm^3^. During the imaging, the transducer array was facing up inside a temperature-regulated water bath for acoustic coupling. The imaged animal was mounted on top of an optically and acoustically transparent animal holder at the transducer array’s focal zone. The animal holder was raster scanned by a 2D motorized translation stage for an expanded 3D-FOV. The 3D-PAULM system has two imaging modes: reversible-switching photoacoustic tomography (RS-PAT) and ultrasound localization microscopy (ULM).

### Data acquisition sequence

The PAT and US data were acquired sequentially. Due to the relatively small focal zone of the 2D semispherical transducer array, raster scanning of the animals by the 2D motorized translation stage was performed. Typically, a total of 6 scanning positions were used (step size: 3 mm), which results in an extended 3D-FOV as needed. In PAT imaging, the excitation wavelength at 750 nm was used with a volume rate of 10 Hz (**Fig. 2c**). The RF data acquired were immediately saved locally, which took approximately 7 ms.

The US data was acquired after the PAT scanning was completed. The synthetic aperture imaging with a 31-element transmission was applied, and the back-scattered echo signals from each transmission event were received with the full aperture, i.e., 256 elements (**Fig. 2c**). A total of 4800 3D US frames were acquired at each scanning position for ULM imaging, with a total acquisition time of ∼3 min. To minimize reverberation artifacts, each US transmission was separated by 150 *µ*s. The received RF data was temporarily stored in the buffer and batch-transferred at once to ensure a US volume rate of 215 Hz. For both PAT and US imaging, 3D images at each scanning location were first reconstructed by using a 3D delay and sum (DAS) method and then stitched to form the final image for further analysis.

### 3D ULM

The 3D US B-mode imaging was performed by using a synthetic aperture method. The in-phase quadratic-phase (IQ) volumes were reconstructed using 3D-DAS with the same voxel size of 60 µm in all dimensions. To mitigate the motion artifacts due to animal breathing, 2D cross-correlation was computed on the lateral maximum intensity projection (MIP) images, using the averaged MIP frame as the reference frame. The extracted correlation threshold of 99% was applied to remove the moving frames, which corresponded to ∼10% of total frames. From the B-mode images, the ULM processing workflow consists of three major steps to reconstruct the vasculature images, as shown in **Supplementary** Fig. 5.

*Step 1.* Spatial-temporal singular value decomposition (SVD) was applied within each block of 600 B-mode volumes to remove clutter signals and extract microbubble signals. The first 150 largest singular values were removed for each block.

*Step 2.* Power Doppler (PD) volumetric images were formed by integrating the SVD-filtered IQ data. To facilitate microbubble separation, directional filtering and bandpass filtering (20-107 Hz) were used.

*Step 3.* Following the filtering, a radial symmetry-based microbubble localization algorithm was applied to extract the sub-diffraction locations of the microbubbles.

*Step 4.* To track the motion of microbubbles across consecutive volumes, a 3D particle tracking algorithm was applied. We assigned a maximum paring/linking search radius of 10 voxels in our tracking algorithm, which corresponds to a maximum bubble velocity of ∼130 mm/s. The maximum gap closing time window was set to 1 which disabled merging the discontinuous track in essence. For frame-to-frame linking we used the linear motion Kalman filter model with default initial state and applied a maximum linking angle of 45 degrees. The microbubble trajectories were then accumulated to generate the ULM density map. Additionally, a microbubble flow velocity map was calculated based on the trajectories, providing information on blood flows

### 3D RS-PAT

To photoswitch and image RpBphP1 or DrBphP-PCM, we combined two CW semiconductor lasers at 635 nm and 790 nm (Civil Laser) and an optical parametric oscillator (OPO) laser (Surelite, Continuum) pumped by an Nd:YAG laser with a 10 Hz pulse repetition rate. The 750 nm light from the OPO laser was used for both PAT imaging and switching off RpBphP1/DrBphP at the same time, while the 635 nm light from the CW laser was used for switching on RpBphP1/DrBphP. The second 790 nm CW laser was used to accelerate the switching-off process. All three laser beams were combined by a dichroic mirror and delivered through the optical fiber bundle. The light beam diameter on the animal surface was ∼1 cm. The maximum light fluence on the skin of the animal was ∼5 mJ/cm^2^, which is below the American National Standards Institute (ANSI) laser safety limit. At each scanning location, a total of 16 cycles of photoswitching were performed. Each switching cycle included 8-second switching on by the 635 nm CW light and 8-second switching off/photoacoustic imaging by the 790 nm CW light and 750 nm pulsed light. Thus, at each scanning location, it took ∼4 min to finish the RS-PAT imaging sequence.

### Imaging processing in RS-PAT

We applied a Hilbert transform to the reconstructed PAT images to extract the signal envelopes, which induces some blurring in the images. Additionally, a 3 by 3 median filter was applied to the Hilbert-filtered images. The differential images were computed by subtracting the OFF-state images from the ON-state images of each switching cycle and then all the differential images were averaged to obtain the final result. When the differential images (e.g., **Fig. 2d**) were displayed, we applied a threshold of three times the noise level, estimated as the standard deviation of the background signals outside the imaged region. The contrast to noise ratio in RS-PAT was calculated as the ratio of the differential signal and the standard deviation of the background.

### Animal preparation for 3D-PAULM

Female Blvra-/- mice or WT mice were used at 4-5 months old with an average weight of 20-30 grams. The imaging region was shaved, and the mouse’s body temperature was maintained at 37°C throughout the experiment. No microbubbles were injected during PAT imaging, but only for ULM imaging. Gas-filled microbubbles with a diameter of 5- μm (VesselVue, Sonovol) were injected via the mouse ocular vein right before the imaging. The total volume of microbubbles injected was less than 100 µL with a concentration of 7×10^8^ microbubbles/mL.

### AAV injection in mouse liver, spleen, kidney, and brain for 3D RS-PAT

Mice were anesthetized with 5% isoflurane in 40% oxygen balanced with nitrogen for anesthesia induction and then moved to the surgical bench. During surgery, the mice subjects inhaled 1.5% isoflurane through a mask, and their body temperature was maintained at 37°C. In preparation for surgery, the ophthalmic ointment was applied to the eyes and 5 mg/kg of Carptofen and Enrofloxacin were administered subcutaneously. Mice were kept in the prone position for liver injections or the left lateral position for spleen injections. Before surgery, the lower back area or the upper abdomen was shaven before surgery and cleaned with iodine and alcohol for kidney and liver/spleen procedures, respectively. A 1 cm longitudinal skin incision was cut below the rib edge at the location of 1 cm lateral to the midline. After opening the abdominal wall, two small retractors were placed, and the organs were exposed. A 25G needle of 5 μl Hamilton syringe was inserted 5 mm into the organs, and 1 μl of AAV solution was slowly injected. The needle remained in position for 5 minutes and was then carefully removed.

For brain injections, mice were mounted on the stereotaxic frame following anesthetization and the top surface of the head was shaved and cleaned. A small skin incision was cut to expose the skull and bregma. Then, a burr hole was drilled on the skull and 1 μl of AAV solution was slowly injected using a pulled glass needle at the location of the hippocampus (1 mm lateral to the midline, -1.5 mm posterior, and 1.5 mm ventral to bregma), striatum (2 mm lateral to the midline, 0 mm posterior, and 3.5 mm ventral to bregma), and lateral hypothalamic area (0.95 mm lateral to midline, -1.40 mm posterior, and 5.25 mm ventral to bregma). The needle remained in place for 10 minutes after injection and was then carefully removed. Following injection, the burr hole was sealed using sterile bone wax, 0.25% Bupicavaine was administered locally, and the skin incision was closed using 6-0 suture. Following surgery, all mice were returned to their home cage to recover from anesthesia.

### Transgenic DrBphP-PCM-4T1 cell preparation for 3D RS-PAT

Lentivirus encoding DrBphP-PCM was used to transduce 4T1 cells to generate a stable line of photoswitchable transgenic cells, henceforth referred to as the 1151 cell line. After flow cytometry sorting on EGFP positive cells, 1151 cells and WT 4T1 cells were seeded at low density and allowed to grow in DMEM (ThermoFisher) supplemented with 10% FBS and 1% Penicillin-Streptomycin (10,000 U/ml). 1151 cells and wild-type 4T1 cells were cultured at 37°C until confluent. The cells were then lifted from culture plates using Trypsin (10 min 37°C) and resuspended in PBS at a concentration of 1 × 10^7^ cells/mL.

### Injecting 4T1 cells into the mammary fat pad

Mice were anesthetized with 2% isoflurane in 30% oxygen balanced with nitrogen through a face mask and kept in a supine position. The fourth nipple skin on the right side was gently held up using a fine tweezer to form a tent shape, and a 27G needle was inserted into the skin next to the nipple until the needle bevel was in. About 0.1 mL (1 × 10^7^ cells/mL) of DrBphP-4T1 cells suspended in PBS were slowly injected. The tweezer was switched to hold the skin around the needle to keep the hole closed while the needle was withdrawn. Then the contralateral fourth mammary fat pad was injected with wild-type 4T1 using the same procedure. After imaging studies the tumor-bearing mice were sacrificed using a carbon dioxide euthanasia when the tumor was larger than 1 cm in size.

### Wide-field fluorescence imaging for validating 3D-PAULM

A lab-made wide-field fluorescence imaging system used a high-sensitivity InGaAs camera that has a broad detection spectrum from 600 nm to 1.7 μm (Ninox 640 II, Raptor Photonics). The camera has a pixel size of 15 μm and a maximum frame rate of 120 Hz. The camera was equipped with an objective lens with an F-number of 1.8 (RPL-OESWIRECON14mmx1.8A, Raptor Photonics). The excitation light source was a 665 nm laser diode (M660L4, Thorlabs). The surface exciton light intensity was ∼50 mW/cm^2^ for all fluorescence imaging in this study. The camera exposure time was 20 ms for all imaging, if not stated otherwise. The fluorescence emission filter was 750/40 nm (FB750-40, Thorlabs) for all imaging. A dichromic mirror with the cut-off wavelength at 730 nm (DMLP730B, Thorlabs) was used.

### Transcranial brain window

To optimize the *in vivo* performance of the two-photon microscopy, we developed a whole-cortex transcranial brain window that was both optically and acoustically transparent. The window was trapezoidal-shaped, with a 9 mm by 10 mm frame size and a useable opening of 8 mm by 7.5 mm. To avoid compressing brain tissue and damaging blood vessels, the window frame had a curved surface that followed the mouse skull’s natural curvature. The window frame was 3D printed using PLA. The window opening was sealed with PVC membrane (10 μm thick) that was both optically and acoustically transparent.

To install the brian window, mice were anesthetized by 100 mg/kg ketamine and 10 mg/kg xylazine. After mounting the mouse head into a stereotactic frame, mice were shaved, and the head surface was cleaned using iodine and alcohol. After midline skin incision, the skin was retracted to fully expose the skull from the olfactory tract area to the occipital area and laterally to the border of the temporal muscles. The skull was kept wet using saline, and two coronal lines at the level of AP −2 mm and AP +4 mm and two sagittal lines along the border of temporal muscles were drilled until the skull became moveable. Using bone wax to seal tiny bleeding sites, the skull was carefully lifted and removed. The transcranial window was then mounted on the skull and glued using Cyanoacrylate (BSI, Atascadero, CA). Next, the skin was glued along the side edges of the window frame. The mouse was returned to the home cage and treated with three days of 5 mg/kg carprofen via subcutaneous (s.c.) injection (Levafen injection, Patterson Veterinary) and seven days of 5 mg/kg enrofloxacin via s.c. injection (Sigma). After a recovery period of seven days, mice were ready for two-photon microscopy imaging.

### Two-photon fluorescence microscopy in the mouse brain

Mice were placed on an imaging stage, stabilized by 3D-printed clips with eye ointment, and anesthetized with isoflurane. *In vivo* imaging to miRFP720 with AAV was performed using a Leica SP8 two-photon DIVE, two weeks after the AAV injection. The miRFP720 excitation wavelength was 1280 nm and the emission filter was 720±25 nm. A 25×/1.05 NA water immersion objective with a motorized correction collar was used to image the mice. The scanning step size was 400 nm along the x and y axis, and 10 µm along the z axis.

## Data Availability

All data supporting the findings of this study are available within the article and Supplementary Information. The major plasmids constructed in this study will be deposited to Addgene.

## Code Availability

The main image processing code used in this study is available at Duke Photoacoustic Imaging Lab’s GitLab page: https://gitlab.oit.duke.edu/pilab/

## ACKNOWLEDGEMENTS

We thank K.Terai and M.Matsuda (Kyoto University, Japan) for donating the Blvra-/- founder mice for establishing our colonies of various transgenic animals, H. Ai (University of Virginia) for the bacterial pBAD-AkaLuc plasmid. Several figures were created with BioRender.com. This work was supported by the grants GM122567 (to V.V.V.), EB028143 (to J.Y.), NS111039 (to J.Y.) and NS115581 (to J.Y. and V.V.V.) from the National Institutes of Health, and CAREER award 2144788 from the National Science Foundation (to J.Y.).

## Author Contributions

L.A.K. generated loxP-BphP1 Blvra-/- homozygous mouse, isolated primary cell cultures, performed the optogenetic experiments, and characterized the miRFP720 in cells. C.M. performed the 3D-PAULM imaging and data analysis, bred the transgenic mice, and assisted with the 3D-PAULM imaging system construction. M.B. performed the molecular cloning of all plasmids and purified the viral particles. H.S. performed animal surgeries for all the 3D-PAULM experiments. Y.T. and T.V. constructed the 3D-PAULM imaging system and assisted with the experiments and data analysis. Y.X. assisted with the 3D-PAULM imaging and data analysis, as well as the animal breeding. L.M. contributed to the image reconstruction algorithm for 3D-PAULM. M.L. and P.S. processed the ULM data for all animal expriments. L.H. performed the 4T1 cell culturing and assisted with developing various AAVs. L.A.K. and V.V.V. designed the optogenetic experiments. C.M. and J.Y. designed the 3D-PAULM imaging experiments. J.Y and V.V.V. supervised the whole project. L.A.K., J.Y. and V.V.V. wrote the manuscript. All authors reviewed and revised the manuscript.

