## Supplementary information for "Advanced deep-tissue imaging and manipulation enabled by biliverdin reductase knockout"

Supplemental Information

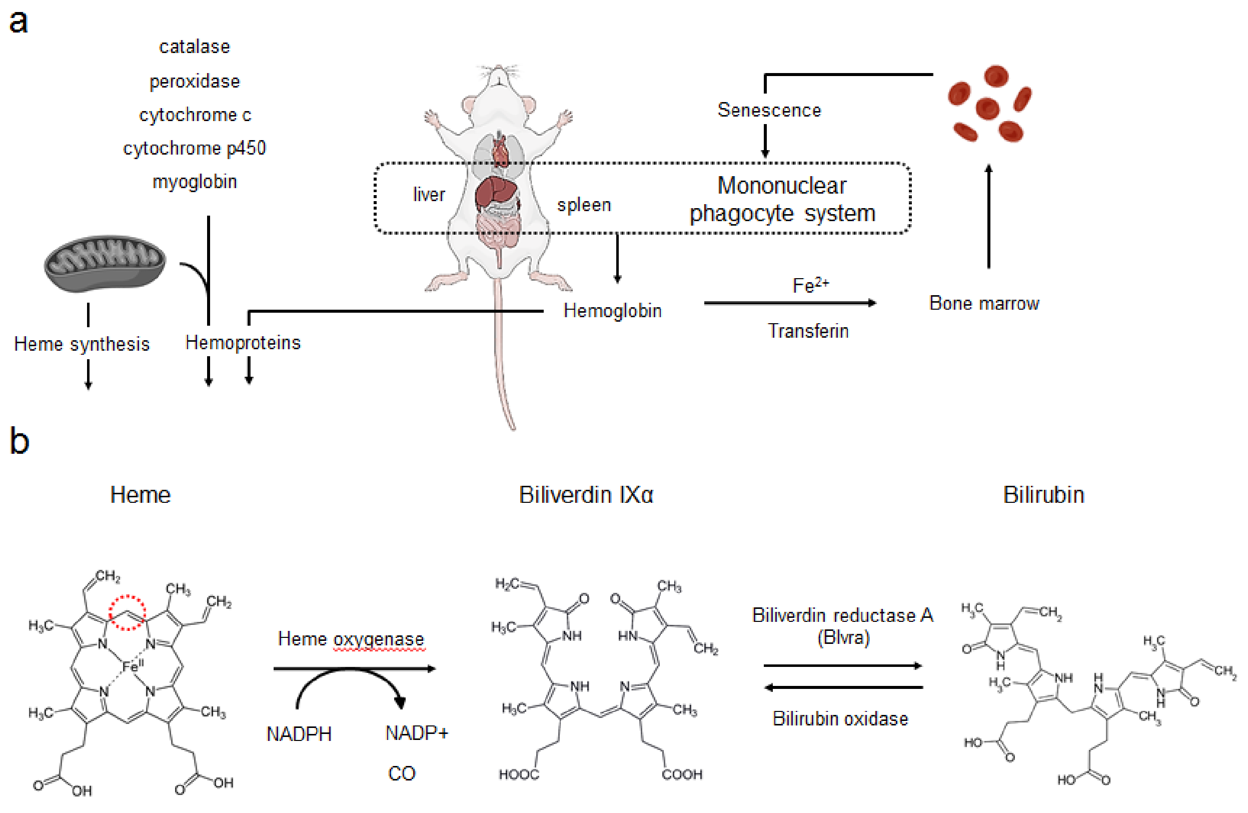


**Supplementary Figure 1. Schematics describing heme metabolism. (a)** Major sites of heme degradation *in vivo*. **(b)** Two steps of heme enzymatic conversion. Heme oxygenase (HO) cleaves the heme ring at the α-methane bridge (indicated with red ring) using the electrons supplied from NADPH. Molecular oxygen enters the reaction resulting in carbon monoxide (CO) production, and the iron is released to be sequestered by ferritin. Biliverdin IXα (BV) can be further converted into bilirubin with BV reductase.

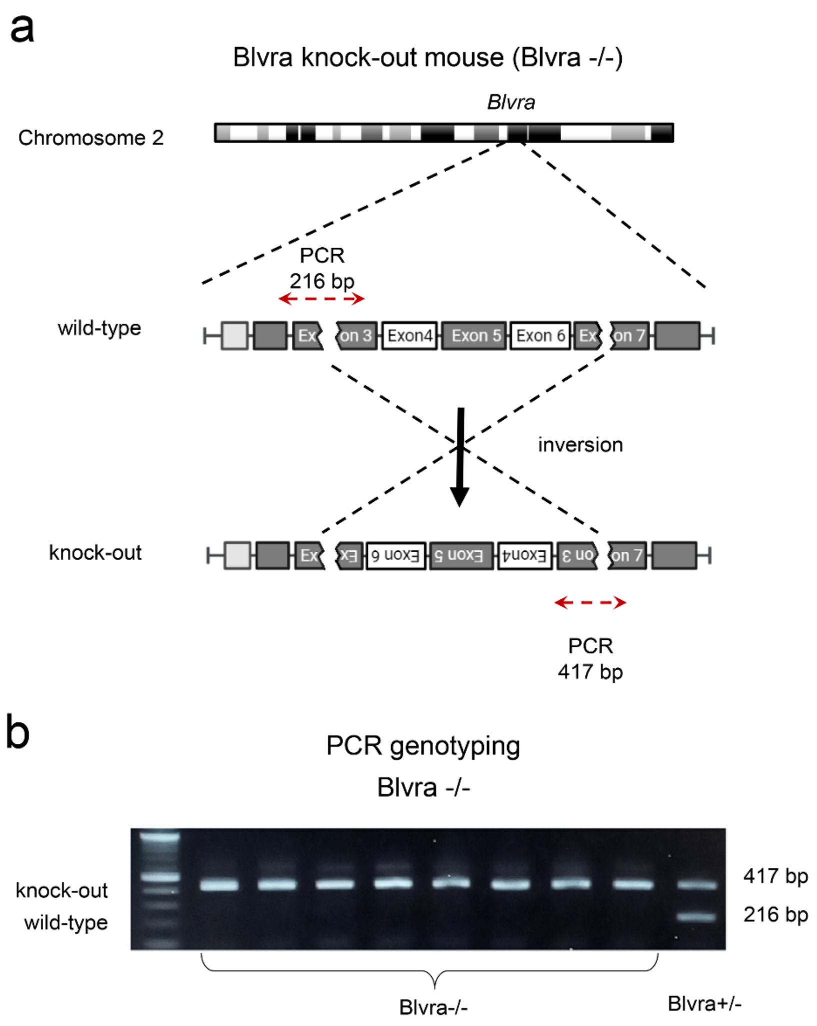


**Supplementary Figure 2. Characterization of Blvra-/- knock-out mouse model. (a)** Schematics of the *Blvra* gene knock-out. Red arrows indicate the regions for primer annealing during PCR genotyping. **(b)** PCR genotyping of Blvra-/- mice. Blvra-/- mice were genotyped using primers annealing to wild-type allele (5'-TGGTAGTGGTTGGTGTTGGC and 5'-CCACTACTCGGCATGGTTCT, amplicon size 216 bp) and to the inverted *Blvra* allele (5'-CCACTACTCGGCATGGTTCT and 5'-AAAGGATGAAAGGCAACATGAGCAG, amplicon size 417 bp).

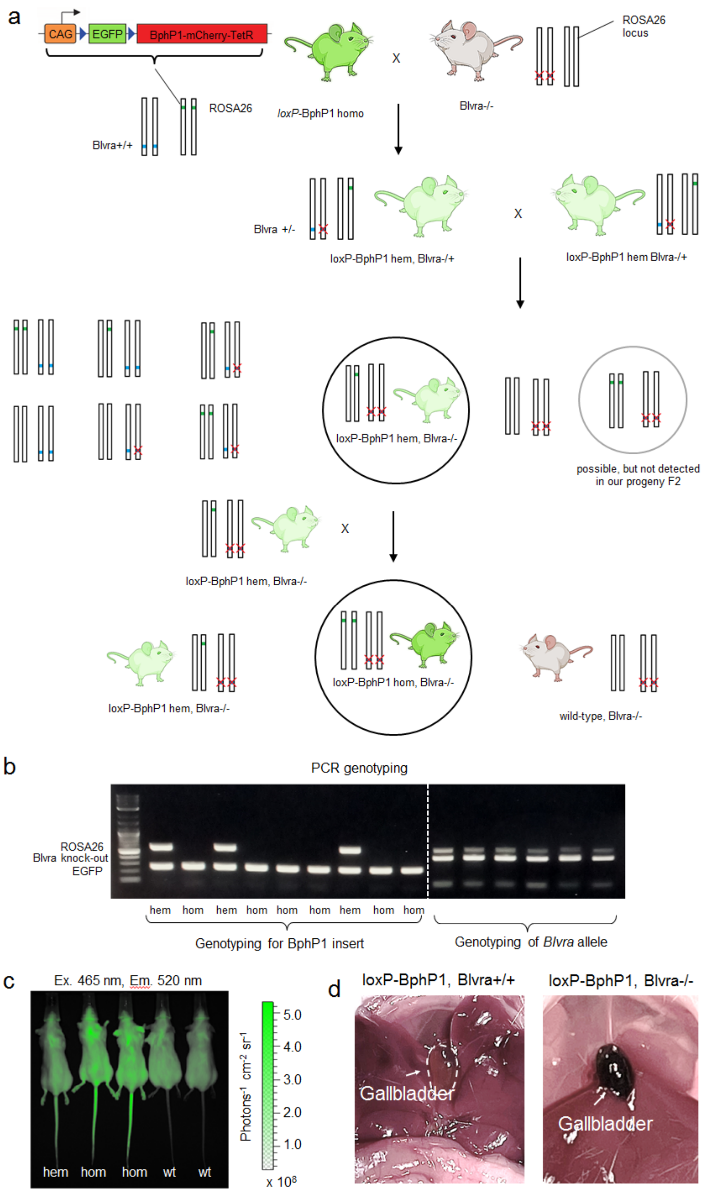


**Supplementary Figure 3. Generation of loxP-BphP1 homo, Blvra-/- mice.** (**a**) Sequential breeding steps for Blvra-/- and loxP-BphP1 mice. **(b)** PCR genotyping of *loxP*-BphP1 homo, Blvra-/- mice was performed using primers annealing to intact *ROSA26* locus (5’-AGCACTTGCTCTCCCAAAGTC, 5’-TGCTTACATAGTCTAACTCGCGAC, product size 564 bp) and transgene floxed *EGFP* sequence (5’-GGCAGAGGATCGTTTCGCGG, 5’-GAAGCACTGCACGCCGTAGG, product size 240 bp). **(c)** *In vivo* fluorescence imaging of progeny to reconfirm *loxP*-BphP1 homo, Blvra-/- genotype. Imaging was performed on the IVIS Spectrum (PerkinElmer). Throughout *in vivo* imaging session animals were maintained under anesthesia with 1.5% vaporized isoflurane. The instrument stage was temperature-controlled (37°C) to avoid hypothermia. For EGFP imaging, 465/20 nm excitation and 520/30 nm emission filters were used. Data were analyzed using Living Image v.4.3 software (Perkin Elmer). **(d)** White-light photographs of the WT and loxP-Bphp1 Blvra-/- mice showing the greenish gallbladder of the loxP-BphP1 Blvra-/- mouse, likely due to the elevated BV level.

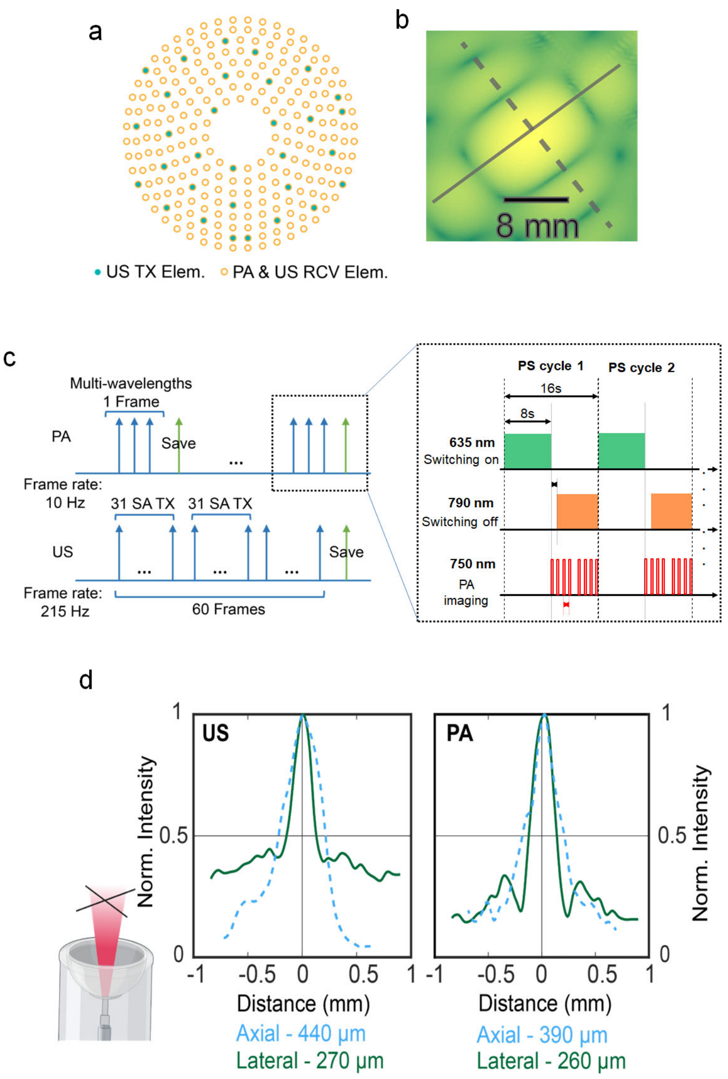


**Supplementary Figure 4. Characteristics of the 3D-PAULM system. (a)** Illustration of the 256-channel 2D semispherical ultrasound transducer array. The 31 transmitting elements of the array (TX) are marked in blue and all the channels are used for receiving both PA and US imaging. **(b)** The simulated acoustic field pressure of the 2D semispherical array using k-wave, showing a 3D field of view of 8 mm in diameter. **(c)** Data acquisition sequence of 3D-PAULM at RS-PAT and ULM modes. RS-PAT uses two continuous-wave lasers at 650 nm and 790 nm for photoswitching the BphP-based probes, and a pulsed laser at 750 nm for photoacoustic imaging of the probes, with a 3D frame rate of 10 Hz. A total of 16 switching cycles were performed at each location with 16 s per cycle. ULM uses a synthetic aperture (SA) based method. A total of 31 transmitting (TX) events are used per image, with a 3D frame rate of 215 Hz. **(d)** The experimentally measured diffraction-limited spatial resolution of the 3D-PAULM system on a cross-hair phantom, showing comparable lateral and axial resolutions for both PA and US imaging when limited by the acoustic diffraction. The lateral and axial resolutions for US are ~440 μm and 270 μm, respectively. The lateral and axial resolutions for PA are ~390 μm and 260 μm, respectively. Please note that the sub-diffraction resolution achieved by the 3D-ULM is demonstrated in the following Supplementary Figure 5 and 6.

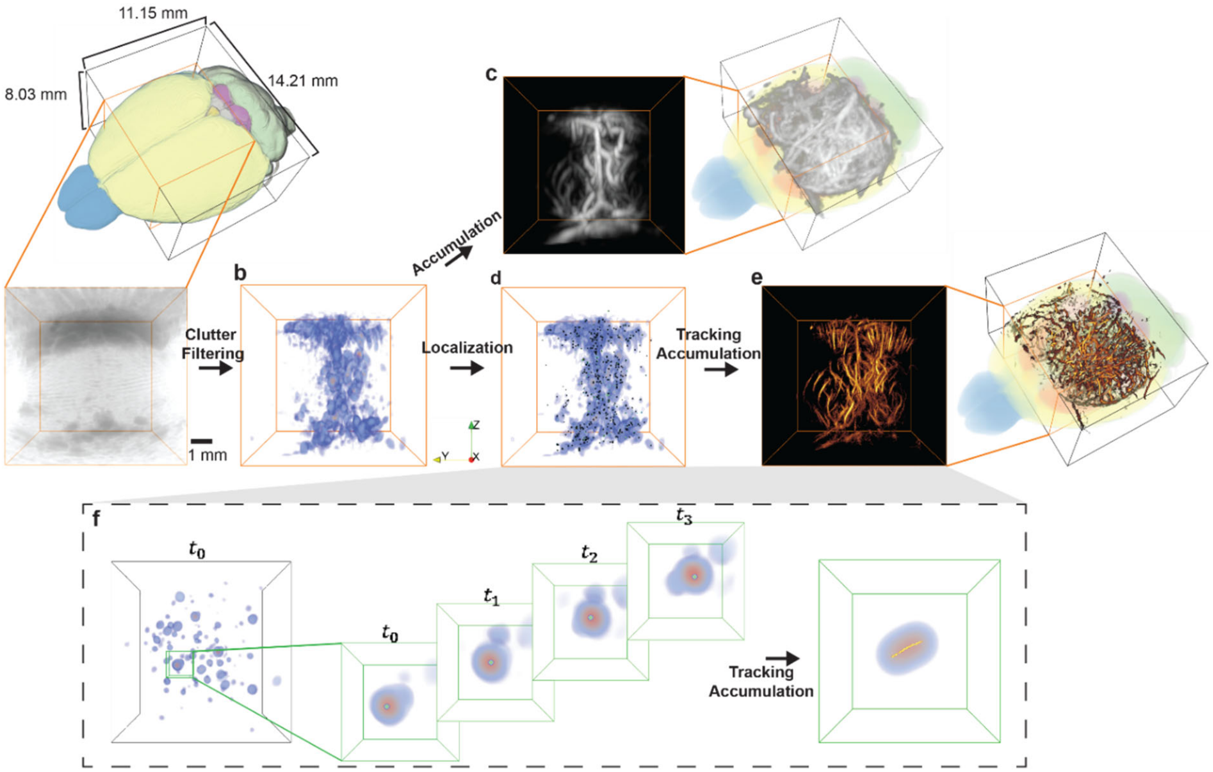


**Supplementary Figure 5. 3D-ULM processing workflows. (a)** US B-mode images are reconstructed by 3D DAS. **(b)** Spatial-temporal singular value decomposition (SVD) filtering is applied to remove the clutter signals. **(c)** Power Doppler image is formed by accumulating the filtered data. **(d)** Radial symmetry-based localization is performed to extract the positions of microbubble signals (green dots). **(e)** Trajectories retrieved using the 3D tracking algorithm are accumulated to form the ULM density map and flow speed map. **(f)** Illustration of MB localization and tracking.

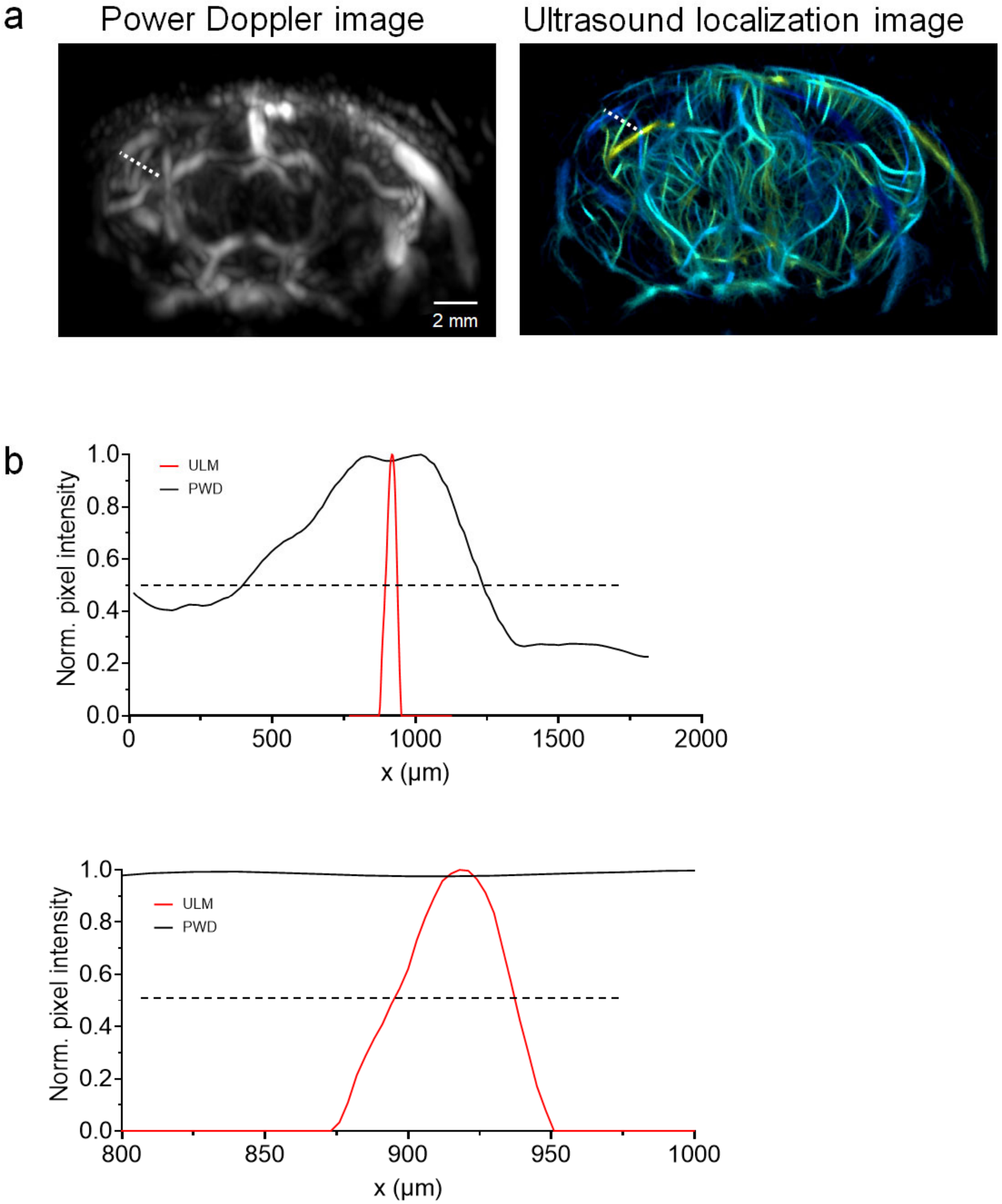


**Supplementary Figure 6. Comparison of ultrasound power Doppler image (PWD) and ultrasound localization microscopy (ULM). (a)** PWD and ULM images of the same mouse brain vasculature. **(b)** The signal profile across a representative vessel marked by the dashed line in **(a)**, showing the full-width at half maximum of 45 μm for ULM and 586 μm for PWD, more than 10-fold resolution improvement.

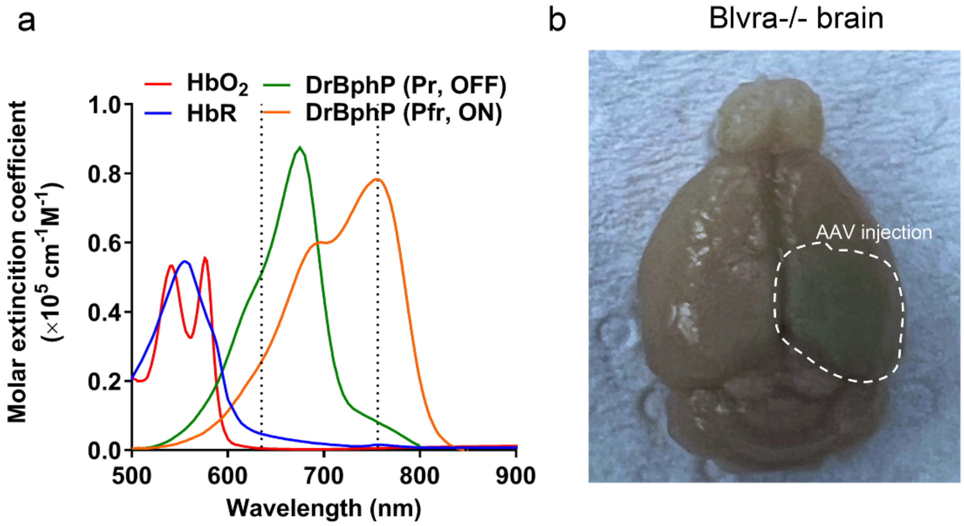


**Supplementary Figure 7. DrBphP-PCM expression in the Blvra-/- brain. (a)** Optical absorption spectra of oxy-hemoglobin, de-oxy-hemoglobin, and DrBphP at Pf (OFF) and Pfr (ON) states. **(b)** White-light photograph of the Blvra-/- brain with the visible green DrBphP expression pattern.

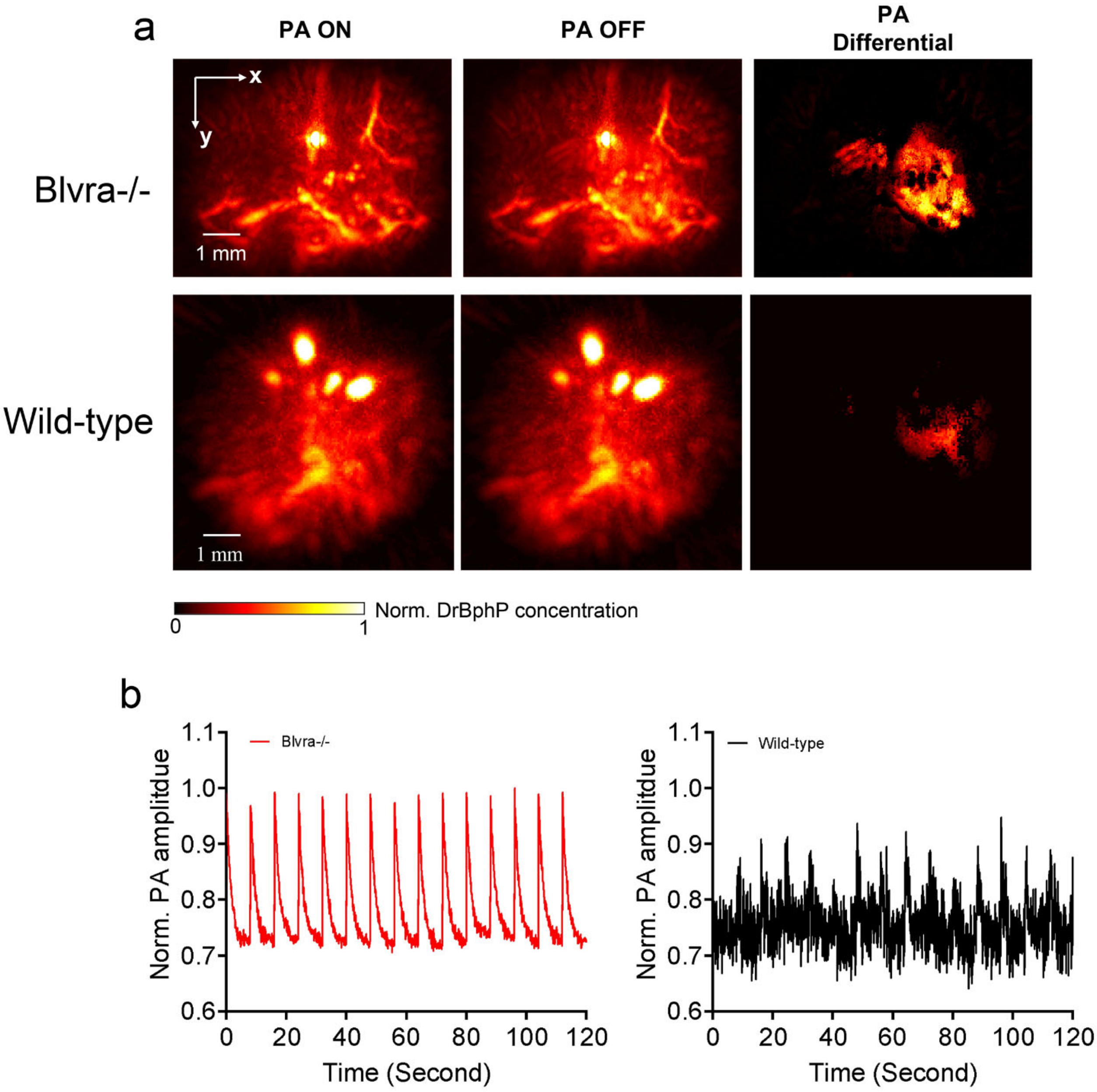


**Supplementary Figure 8. RS-PAT of the DrBphP-PCM expressed in the Blvra-/- and WT brains.** **(a)** *x-y* projection PA images at ON- and OFF-state, as well as the differential image, in the Blvra-/- and WT brains, showing much stronger DrBphP-PCM signals in the Blvra-/- brain. **(b)** Photoswitching dynamics of the PA signals in the Blvra-/- and WT brains, showing a much stronger and cleaner signal switching in the Blvra-/- brain.

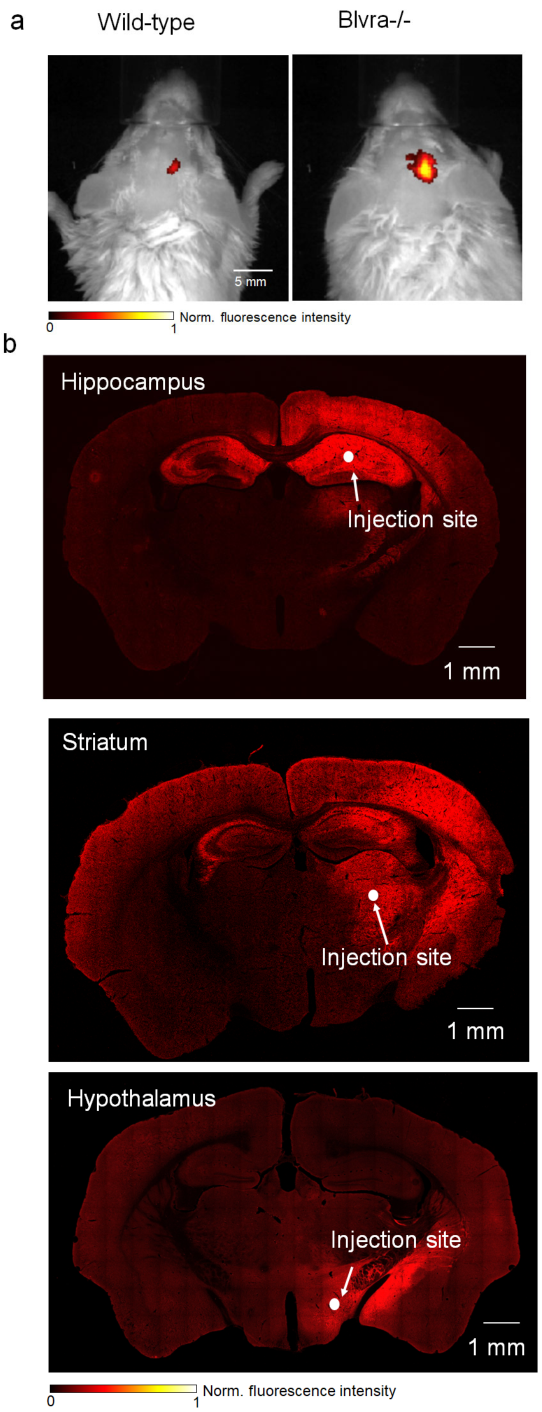


**Supplementary Figure 9. Fluorescence image of the DrBphP-PCM expression in deep mouse brain. (a)** Wide-field fluorescence image of the DrBphP-PCM expression in the Blvra-/- mouse and the wild-type. **(b)** Confocal fluorescence images of DrBphP-PCM expressed in hippocampus, striatum, and hypothalamus regions of the Blvra-/- brain. The brain was perfused and sliced with a 5 µm thickness. The approximate injection sites are marked by white arrows. The signal breaching into other brain regions is clearly visible in the hippocampus and striatum injections.

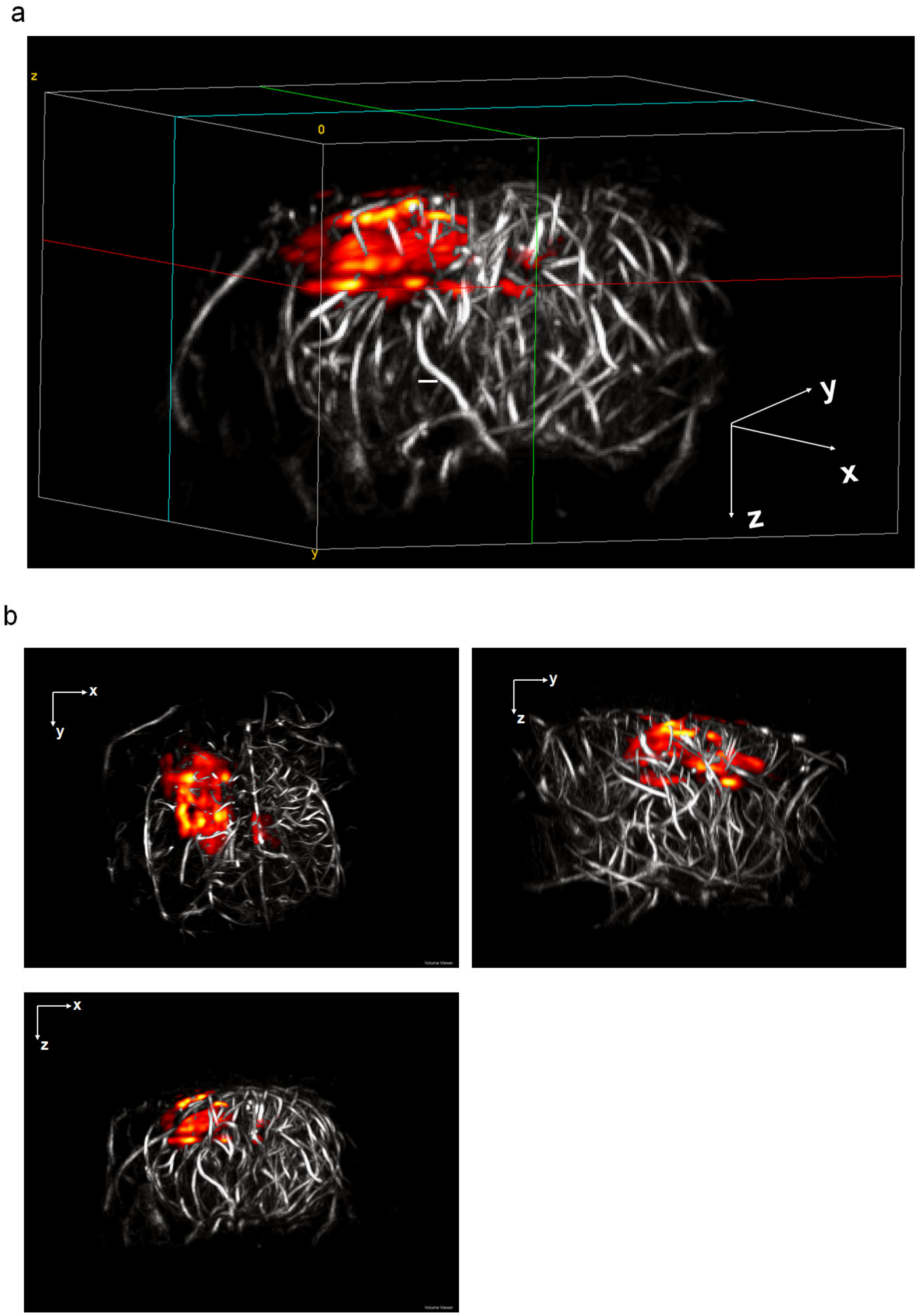


**Supplementary Figure 10. 3D-PAULM volumetric rendering of the DrBphP-PCM expressed in hippocampus of the Blvra-/- brain.** The ULM images of the brain vasculature are shown in gray and the RS-PAT images of the DrBphP-PCM-expressing cells are shown in color. The imaging processing is performed with VolView (Kitware).

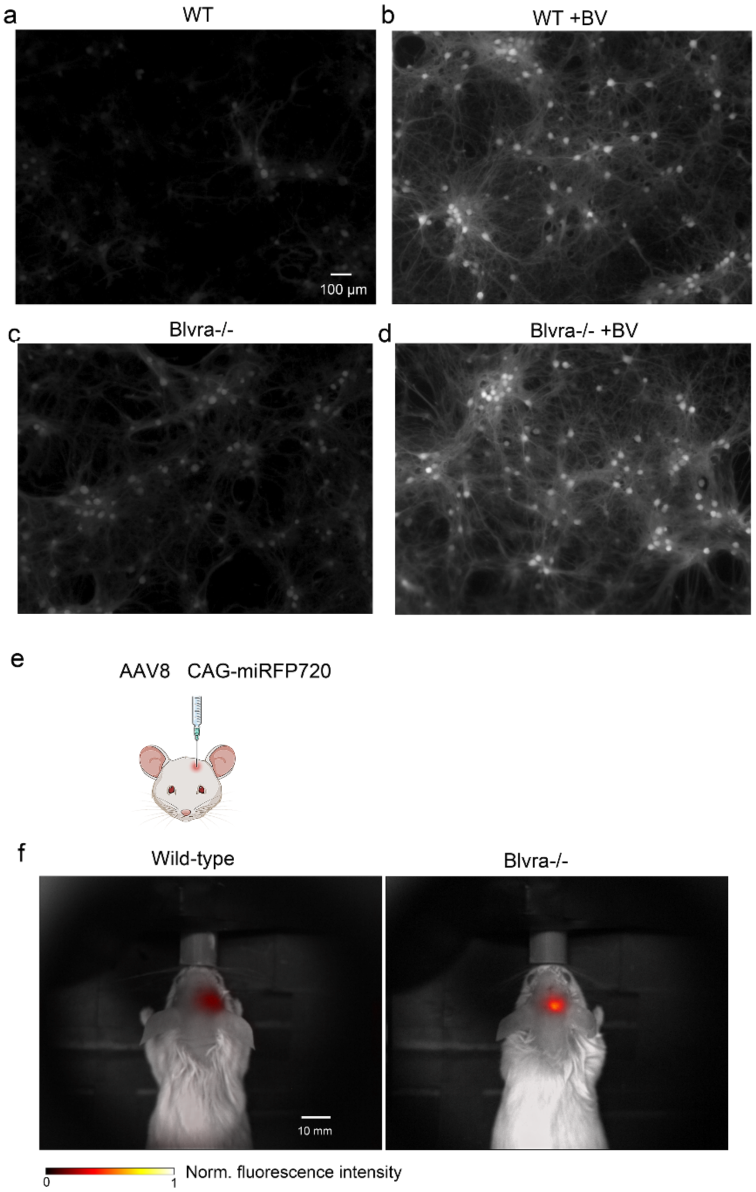


**Supplementary Figure 11. Fluorescence microscopy of primary cortical neurons and live brain transduced with AAV8 CAG-miRFP720. (a-d)** For fluorescence microscopy, neurons were plated on poly-D-lysine pre-coated MatTek glass-bottom dishes (35 mm). Cells were transduced with AAV8 CAG-miRFP720 (multiplicity of infection of $10^5$) on 5 DIV and cultured in Neurobasal Plus Medium with B-27 Plus Supplement (Gibco), 1 mM GlutaMAX (Gibco, 35050061), 100 U/ml penicillin and 100 μg/ml streptomycin for 12 days before imaging. Where necessary cells were supplemented with 2 μM BV 24 h before imaging. Fluorescence of miRFP720 was recorded in Cy5.5 channel (Ex. 665/45 nm, Em. 725/50 nm) for WT (a, b) and Blvra-/- (c, d) cells using the same acquisition settings. The Olympus IX81 microscope was operated with a SlideBook v.6.0.8 software (Intelligent Imaging Innovations). (**e-g**) *In vivo* wide-field fluorescence imaging of mouse brain transduced with AAV8 CAG-miRFP720 by local injection in the dorsal hippocampus (surface to injection site: ~2-3 mm) of the right hemisphere. The fluorescence signals were recorded at an excitation of 650 nm and an emission of 750 nm.

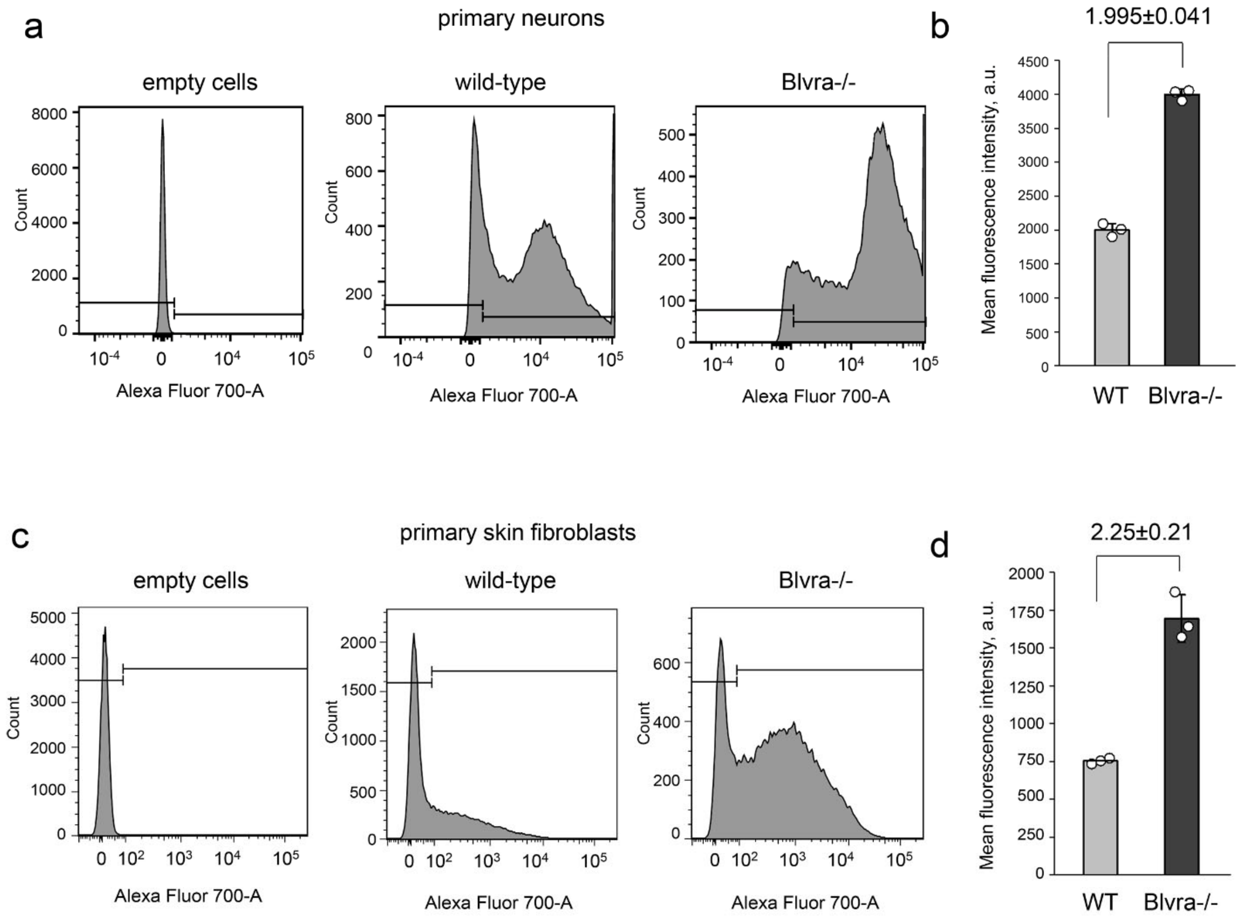


**Supplementary Figure 12. Flow cytometry analysis of primary neurons and skin fibroblasts expression miRFP720. (a, b)** Primary WT and Blvra-/- cortical neurons were transduced with AAV8 CAG-miRFP720 (multiplicity of infection of $10^5$) on 5 DIV and cultured in Neurobasal Plus Medium with B-27 Plus Supplement (Gibco), additional 1 mM GlutaMAX (Gibco, 35050061), 100 U/ml penicillin and 100 µg/ml streptomycin for 12 days before analysis. Gating of miRFP720 positive cortical neurons (a) and quantitative difference in mean fluorescence intensity (b) are shown. **(c, d)** Primary WT and Blvra-/- skin fibroblasts were transduced with AAV8 CAG-miRFP720 (multiplicity of infection of $10^5$) on 5 DIV and cultured in DMEM, 10% FBS, 100 U/ml penicillin and 100 µg/ml streptomycin for 12 days before analysis. Gating of miRFP720 positive cells (c) and quantitative difference in mean fluorescence intensity (d) are shown. Data are presented as mean values +/- SD (*n*=3; independent transfection experiments).

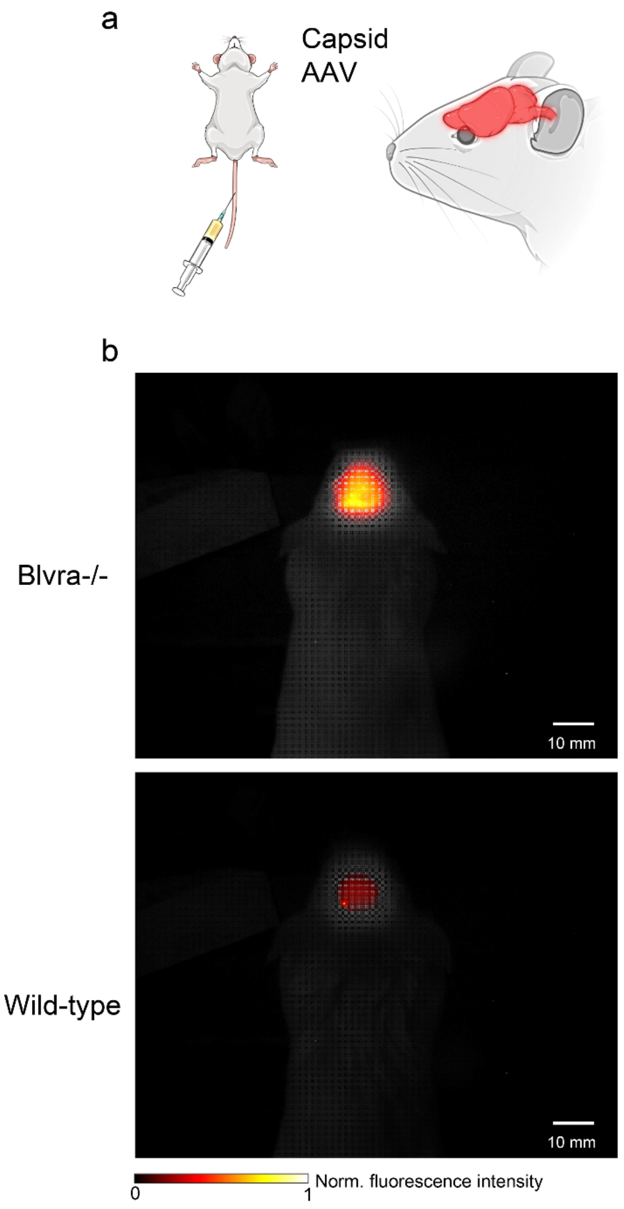


**Supplementary Figure 13. Whole-brain expression of DrBphP-PCM using the blood-brain penetrating AAV-PHP.eB capsids. (a)** Illustration of the whole-brain expression of DrBphP-PCM via tail injection of AAV-PHP.eB DrBphP-PCM. (**b**) Wide-field DrBphP-PCM fluorescent images of the Blvra-/- and WT brain *in vivo*, showing a 2-fold increase of the fluorescence intensity in the Blvra-/- brain. The DrBphP signals are relatively uniform across the whole brain for both Blvra-/- and WT mice.

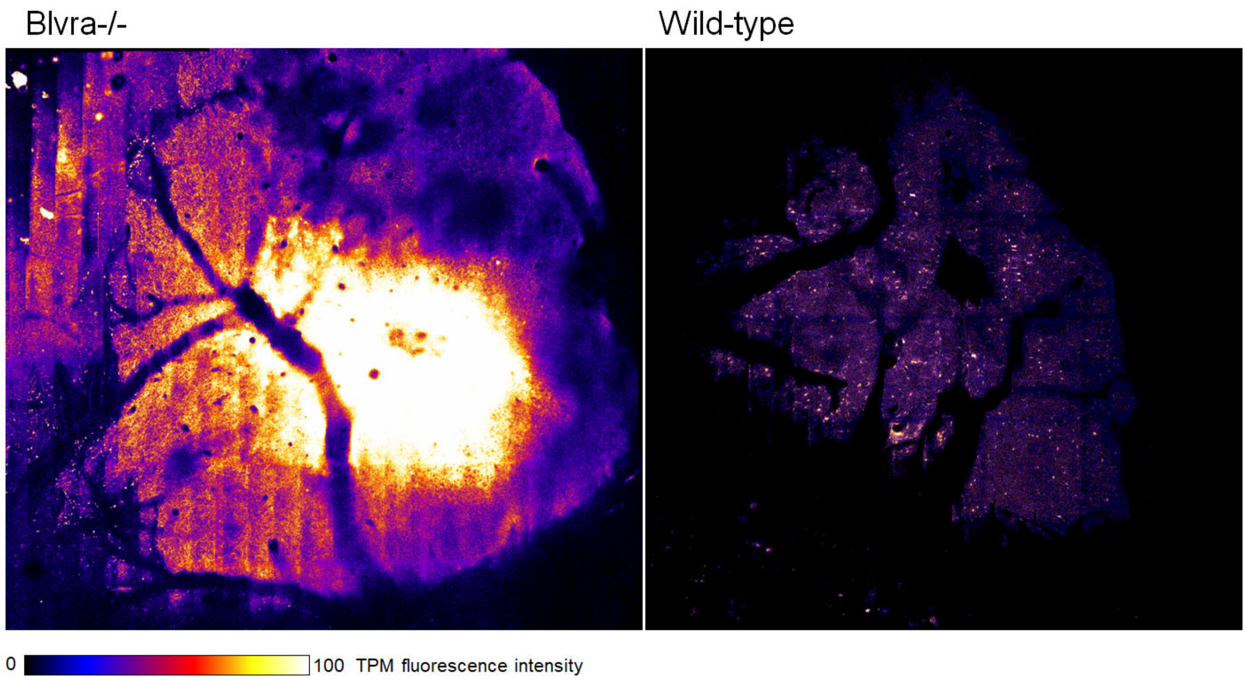


**Supplementary Figure 14. Two-photon microscopy of the AAV-mediated miRFP720 expression in the brain of (a) Blvra-/- mouse and (b) wild-type mouse.** The AAV injection was at 0.8 mm beneath the brain surface.

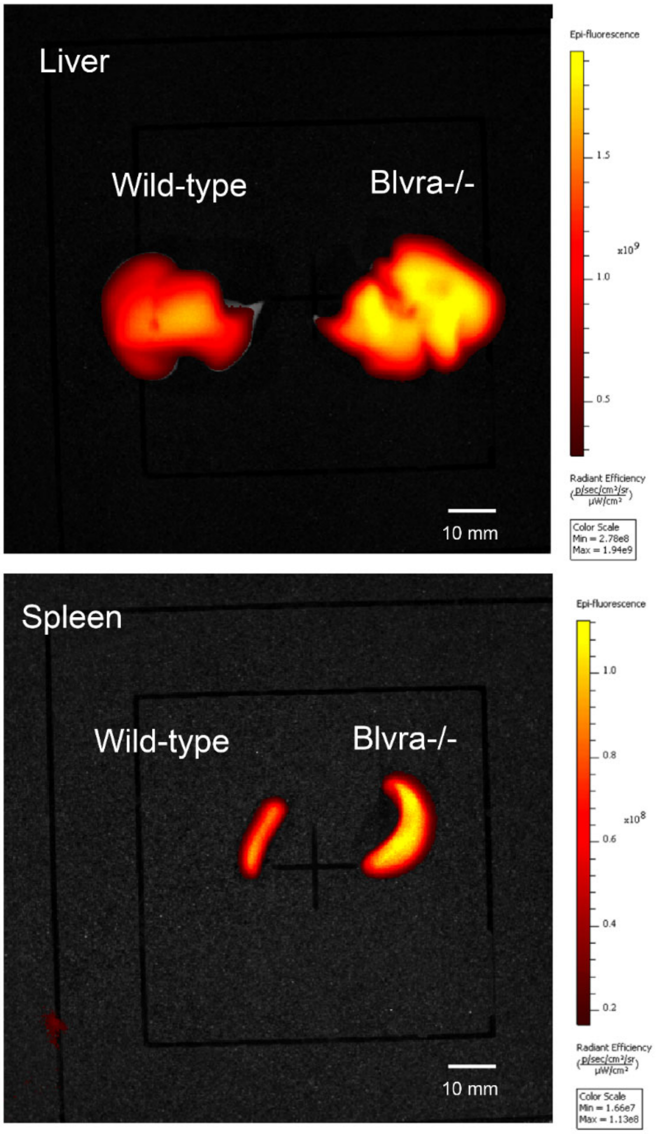


**Supplementary Figure 15. Fluorescence images of the extracted liver and spleen of loxP-BphP1, Blvra-/- and in loxP-BphP, Blvra+/+ mice.** AAV-Cre-mediated expression of the genomically encoded BphP1 in the liver and spleen are both stronger in the Blvra-/- mouse than the WT (Blvra+/+) mouse. Images were acquired with IVIS Spectrum (Perkin Elmer).

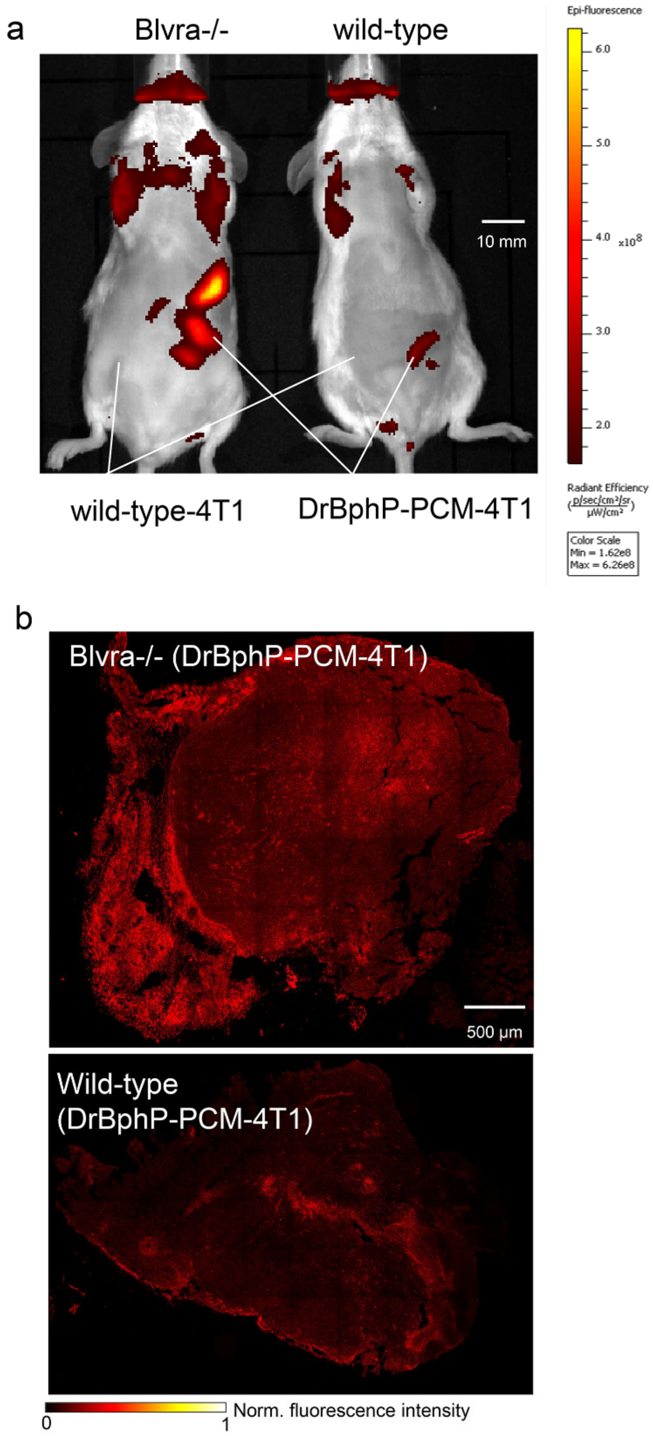


**Supplementary Figure 16. Fluorescence imaging of the implanted 4T1 tumors in Blvra-/- and WT mice. (a)** IVIS fluorescent images of the tumor-bearing mice *in vivo*, showing that the DrBphP-PCM-4T1 tumor in the Blvra-/- mouse has the strongest signals. **(b)** Confocal microscopy images of the extracted and sliced DrBphP-PCM-4T1 tumors from the Blvra-/- and WT mice, showing that the Blvra-/- mouse has a relatively uniform signal intensity throughout the tumor.
